# Maf1, a repressor of RNA polymerase III-dependent transcription, regulates bone mass

**DOI:** 10.1101/2021.11.09.467887

**Authors:** Ellen Busschers, Naseer Ahmad, Li Sun, James Iben, Christopher J. Walkey, Aleksandra Rusin, Tony Yuen, Clifford J. Rosen, Ian M. Willis, Mone Zaidi, Deborah L. Johnson

**Affiliations:** Department of Molecular and Cellular Biology, Baylor College of Medicine, Houston, TX 77030, USA; Departments of Medicine and Pharmacological Sciences and Center for translational Medicine and Pharmacology, Icahn School of Medicine at Mount Sinai, New York, NY 10029, USA; Molecular Genomics Core, Eunice Kennedy Shriver National Institute of Child Health and Human Development, National Institutes of Health, Bethesda, MD 20892 USA; Center for Clinical and Translational Research, Maine Medical Center Research Institute, Scarborough, Maine 04074; Departments of Biochemistry and Systems and Computational Biology, Albert Einstein College of Medicine, Bronx, NY 10461, USA

## Abstract

Maf1, a key repressor of RNA polymerase III-mediated transcription, has been shown to promote mesoderm formation *in vitro*. Here, we show, for the first time, that Maf1 plays a critical role in the regulation of osteoblast differentiation and bone mass. A high bone mass phenotype was noted in mice with global deletion of *Maf1* (*Maf1*^*-/-*^ mice), as well as paradoxically, in mice that overexpressed MAF1 in cells of the osteoblast lineage (*Prx1*-Cre;LSL-*Maf1* mice). Osteoblasts isolated from *Maf1*^*-/-*^ mice showed reduced osteoblastogenesis *ex vivo. Prx1*-Cre;LSL-*MAF1* mice showed enhanced osteoblastogenesis concordant with their high bone mass phenotype. Thus, the high bone mass phenotype in *Maf1*^*-/-*^ mice is likely due to the confounding effects of the global absence of Maf1 in *Maf1*^*-/-*^ mice. Expectedly, MAF1 overexpression promoted osteoblast differentiation and shRNA-mediated Maf1 downregulation inhibited differentiation of ST2 cells, overall indicating Maf1 enhances osteoblast formation. We also found that, in contrast to MAF1 overexpression, other perturbations that repress RNA pol III transcription, including Brf1 knockdown and the chemical inhibition of RNA pol III by ML-60218, inhibited osteoblast differentiation. All perturbations that decrease RNA pol III transcription, however, enhanced adipogenesis in ST2 cell cultures. RNA-seq was used to determine the basis for these opposing actions on osteoblast differentiation. The modalities used to alter RNA pol III transcription resulted in distinct changes gene expression, indicating that this transcription process is highly sensitive to perturbations and triggers diverse gene expression programs and phenotypic outcomes. Specifically, Maf1 induced genes in ST2 cells known to promote osteoblast differentiation. Furthermore, genes that are induced during osteoblast differentiation displayed codon bias. Together, these results reveal a novel role for Maf1 and RNA pol III-mediated transcription in osteoblast fate determination and differentiation and bone mass regulation.

## Introduction

RNA polymerase (pol) III transcribes various untranslated RNAs including 5S rRNA and tRNAs. RNA pol III-derived transcripts play essential roles in several processes, including protein synthesis and secretion (1, 2). In addition to RNA pol III, transcription of tRNAs requires the recruitment of TFIIIC and TFIIIB to the promoter. TFIIIB consists of Brf1, TATA binding protein (TBP) and B double prime (Bdp1) (3). RNA pol III-dependent transcription is also tightly regulated through either direct or indirect mechanisms that control TFIIIB recruitment to the promotor (4–11). Maf1 is a key repressor of RNA pol III-dependent transcription (12, 13). It binds to RNA pol III and prevents the interaction between RNA pol III and TFIIIB (14, 15). Maf1 has also been shown to regulate a variety of RNA pol II-transcribed targets (13, 16–19), and acts as a tumor suppressor (16, 18), regulates metabolism (20, 21), and longevity (22, 23).

RNA pol III-mediated transcription has been shown to play an important role during development and cellular differentiation. Two RNA pol III isoforms are differentially expressed between pluripotent and differentiated embryonic stem cells (ESCs) (24–26). RNA pol III-dependent transcription also modulates the formation of hematopoietic lineages in zebrafish (27) and transcription is downregulated during skeletal muscle differentiation of *Xenopus tropicalis* (28). Maf1 enhances the formation of mesoderm in embryonic stem cells, and the downregulation of RNA pol III-dependent transcription enhances the differentiation of ESCs and 3T3-L1 cells into adipocytes (29).

Diseases associated with ribosomal disfunctions, ribosomopathies, are commonly associated with bone marrow, skeletal and craniofacial disorders (30). This surprising tissue specificity suggests that these tissues may be particularly sensitive to alterations in protein synthesis. For example, Treacher Collins syndrome is caused by mutations in *POLR1C, POLR1D* or *TCOF1*, which affect rDNA transcription by RNA pol I and ribosome biogenesis (31, 32). RNA pol III-derived transcripts may also play a role as POLR1C and POLR1D are common subunits of both RNA pol I and RNA pol III. In yeast, Treacher Collins syndrome related mutations in *POLR1D* result in altered functions of both RNA pol I and III (33). Furthermore, RNA pol III-dependent transcription may play a role in Cerebellofaciodental syndrome which is associated with mutations in the RNA pol III-specific transcription factor, BRF1. This syndrome is characterized by a neurodevelopmental phenotype as well as changes in the facial and dental structure and delayed bone age (34–36). Additionally, bone phenotypes have been described in a subgroup of patients with mutations in RNA pol III subunits (34–38).

It is known that Maf1 regulates mesoderm formation and adipocyte differentiation (29). Osteoblasts and adipocytes are both derived from the mesenchymal lineage (39) and mutations in *BRF1* and RNA pol III subunits are associated with bone-related phenotypes. These facts led us to hypothesize that regulation of RNA pol III-dependent transcription by Maf1 may play a fundamental role in osteoblast differentiation, bone formation and hence, bone mass. Here, we show that both whole body deletion of *Maf1* in mice and tissue specific overexpression of MAF1 in stromal cells of the long bones enhances bone mass *in vivo*, while Maf1 induces osteoblast differentiation *in vitro*. To further examine the role of MAF1 and RNA pol III-mediated transcription in osteoblast differentiation, we attempted to study the effect on osteoblast differentiation by repressing transcription through different approaches. Surprisingly, while Maf1 induced osteoblast differentiation, repression of RNA pol III-dependent transcription, by either chemical inhibition of RNA pol III or by Brf1 knockdown, decreased osteoblast differentiation. Thus, changes in Maf1 expression produce an opposing effect on osteoblast differentiation compared with other approaches that repress RNA pol III-mediated transcription. We further show that these three different approaches to decrease RNA pol III-dependent transcription result in divergent gene expression changes. Altered Maf1 expression affects the expression the osteoblast differentiation gene program. Together, these findings reveal that Maf1 plays a key role in osteoblast differentiation and bone mass regulation and that its ability to regulate RNA pol III-dependent transcription contributes to the observed phenotypic outcomes.

## Results

### Maf1 Overexpression Stimulates Osteoblast Lineage Cells to Differentiate into Mature Osteoblasts, and Enhances Adipogenesis

To determine the role of Maf1 on bone mass and bone formation *in vivo* we examined the bone phenotype of the global *Maf1*^*-/-*^ mouse model (21). Micro computed tomography (µCT) was used to determine femur, tibia and spine bone mineral density (BMD) and microstructural parameters in mature male mice at 12 weeks of age. When compared to their WT littermates, *Maf1*^*-/-*^ mice showed a significant increase in bone volume, trabecular number and trabecular thickness at the spine and increased bone volume and trabecular thickness in the tibia. Femur samples showed a similar trend without reaching statistical significance (Suppl. Fig. 1). To determine changes at the cellular level, we isolated primary bone marrow stromal cells and hematopoietic cells from these mice to determine their capacity *ex vivo* to form osteoblasts and osteoclasts, respectively. Surprisingly, osteoblast formation was reduced, while osteoclastogenesis was increased in cells derived from *Maf1*^*-/-*^ mice (Suppl. Fig. 2). While this suggests that Maf1 affects osteoblast and osteoclast formation in long term *ex vivo* cultures, this change does not reflect the increase in bone volume seen in the *Maf1*^*-/-*^ mice, which is likely the result of non-cell-autonomous effects arising from the deletion of *Maf1* in other tissues. The *ex vivo* data, however, indicate that Maf1 is a positive regulator of osteoblast differentiation, and its overexpression in osteoblasts would therefore result in increased osteoblastogenesis.

We therefore determined the effect of Maf1 overexpression specifically in the long bones by developing a transgenic mouse strain with an HA-tagged *MAF1* construct inserted in the *Rosa 26* locus. The expression of MAF1 was driven by a hybrid cytomegalovirus enhancer chicken β-actin (CAGGS) promoter with a lox-stop-lox cassette inserted between the promoter and *MAF1*. This *Rosa-lox-stop-lox-MAF1-HA* (LSL-MAF1) strain was then crossed to a *Prx1*-Cre mouse to overexpress MAF1 in the mesenchyme of the developing limb bud (40). HA expression in the femur following Cre-recombination was confirmed by western blotting. qRT-PCR showing ∼16-fold increase in MAF1 mRNA expression compared to endogenous Maf1 (Fig. 1A-B). To assess the effect of increased MAF1 expression on bone mass, we employed µCT imaging on femurs of 12-week-old male mice. MAF1 overexpression in *Prx1*-Cre; LSL-MAF1 mice led to an increase in bone volume, trabecular number and thickness, and connectivity density, and a corresponding reduction in trabecular separation when compared with Cre-controls. Cortical thickness was not significantly increased (Fig. 1C).

**Figure 1.**
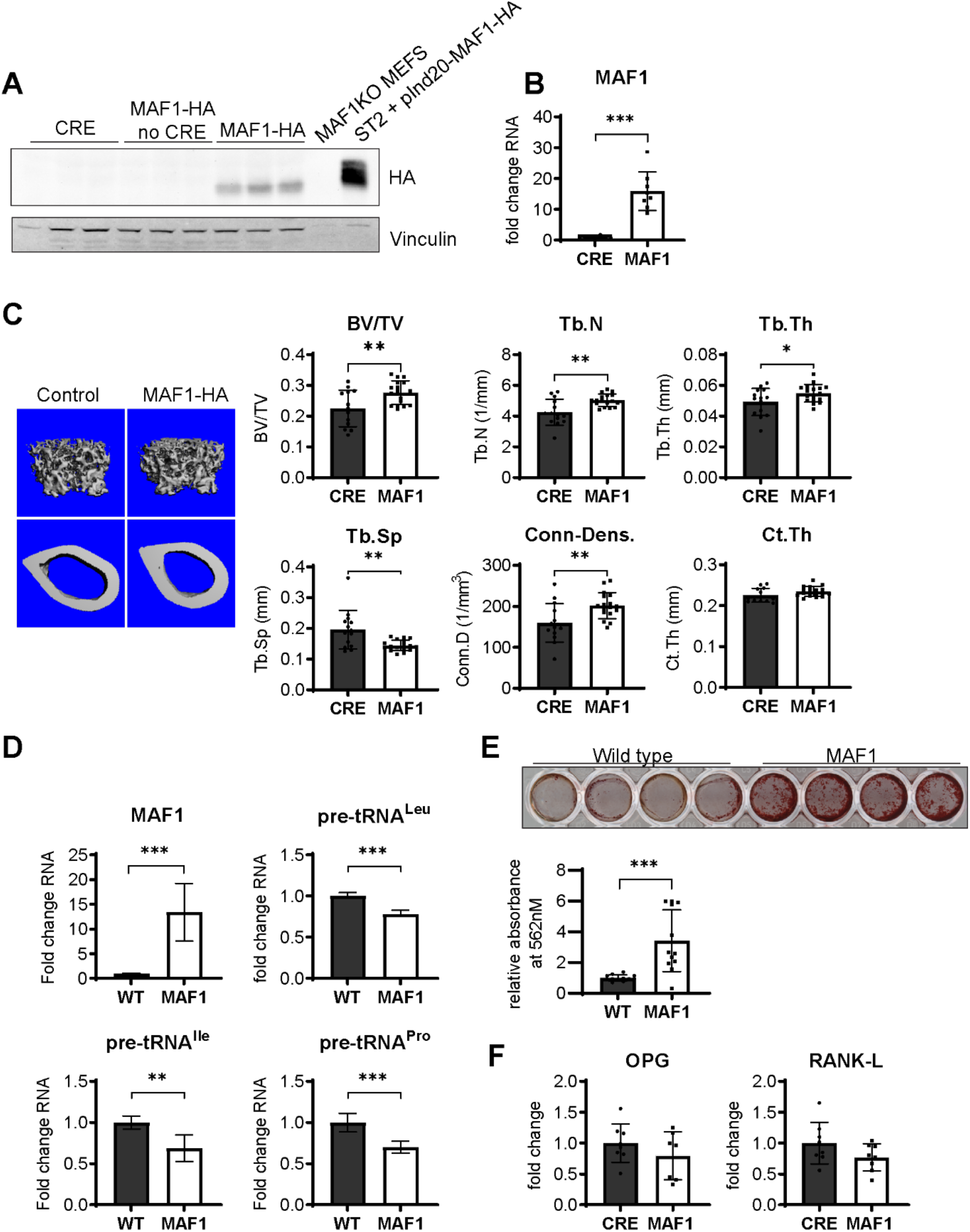
Bone specific overexpression of MAF1-HA increases bone volume in mice. **(A**) Western blot of HA expression in the femur of 12 week old male Prx1-Cre MAF1-HA mice compared to Prx1-Cre - WT and WT-MAF1-HA mice. (**B**) qRT-PCR analysis showing MAF1 RNA in femurs from Prx1-Cre-MAF1 mice and control Prx1-Cre mice. (**C**) Left, representative images of µCT of femoral bone. Right, quantification of µCT analysis: Bone volume/total volume (BV/TV), trabecular number (Tb.N), trabecular thickness (Tb.Th), trabecular separation (Tb.Sp), connectivity density (Conn-Dens.) and cortical thickness (Ct.Th). n= 13 for Prx1-Cre and n=17 for MAF1 mice. (**D**) qRT-PCR of MAF1 and pre-tRNAs in primary stromal cells isolated from 6-8 week old WT or MAF1 overexpressing mice. (**E**) Representative plate of Alizarin red labeled mineralization of WT and MAF1-HA primary stromal cells (top). Quantification of Alizarin red after destaining with 10% CPC. (**F**) qRT-PCR of Opg and Rankl in Prx1-Cre and MAF1 overexpressing femurs at 12 weeks. Results represent means ±SD, *P<0.05, **P<0.01, ***P<0.001 determined by Student’s t test. Figure 1 - Source Data contains uncropped images of western blots.

To determine the effect of MAF1 overexpression at the cellular level, primary stromal cells were isolated from femurs and tibia of *Prx1*-Cre; LSL-MAF1 mice and cultured *ex vivo*. These primary cells displayed an increase in MAF1 expression and showed a corresponding decrease in tRNA gene transcription (Fig. 1D). When Maf1-overexpressing stromal cells were allowed to differentiate into bone-forming osteoblasts in media containing ascorbic acid and β-glycerolphosphate, there was a clear increase in their mineralizing capacity on alizarin red staining (Fig. 1E). These results show that Maf1 triggers the differentiation of osteoblasts into mineralizing cells. We found no change in the pro-osteoclastogenic cytokines, Receptor activator of NF-κβ ligand (RANKL) and osteoprotegerin (OPG) (Fig. 1F).

To further confirm a role for Maf1 on osteoblast differentiation, we overexpressed MAF1 in the mouse stromal cell line ST2 using a dox-inducible MAF1-HA construct. Cells stably expressing either the MAF1-HA construct or a control vector were treated with 1µM doxycycline (Dox) and differentiated into osteoblasts. Ectopic MAF1 expression was confirmed by Western blot (Fig. 2A) and qRT-PCR and resulted in the reduction of pre-tRNA^Ile^ and pre-tRNA^Leu^ gene transcription (Fig. 2B). Maf1 overexpression resulted in enhanced staining for alkaline phosphatase (Alp), a marker for early osteoblast differentiation (Fig. 2C), as well as increased *in vitro* mineralization as noted on alizarin red staining (Fig. 2D). Maf1 overexpression also resulted in a significant increase in expression of osteoblast marker genes, namely *Col1A, Sp7, Osteocalcin, Alp* and bone sialoprotein (*Bsp)* (Fig. 2E).

**Figure 2.**
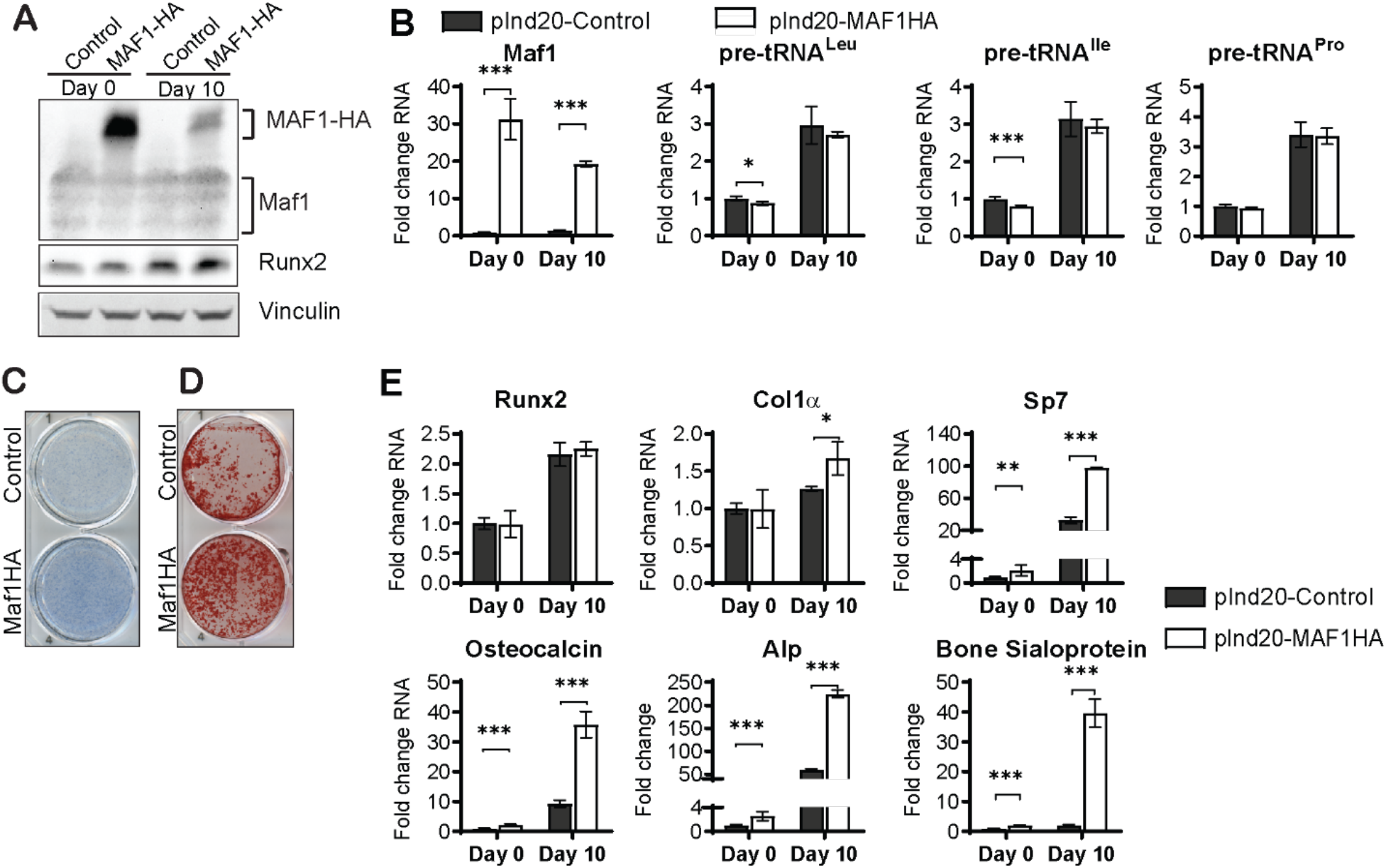
MAF1 increases *in vitro* osteoblast differentiation and mineralization. ST2 cells were infected with a dox-inducible pInd20-MAF1HA or control construct. Cells were treated with 1µM Dox starting 1 day before differentiation was started. (**A**) Western blot analysis showing Maf1, Runx2 and Vinculin in ST2 cells differentiated into osteoblast at day 0 and day 10. (**B**) qRT-PCR analysis showing Maf1 and pre-tRNA expression in ST2 cells pre- and during osteoblast differentiation. (**C**) Representative image of Alp staining of control and MAF1-HA expressing cells. (**D**) Representative image of alizarin red analysis of ST2 cells overexpressing control or MAF1-HA after culture in osteoblast differentiation medium. (**E**) qRT-PCR analysis showing relative expression of Runx2, Col1α, Sp7 (Osterix), Alp and Bone sialoprotein before and 10 days after addition of osteoblast differentiation medium. Results represent means ±SD of three independent replicates, *P<0.05, **P<0.01, ***P<0.001 determined by Student’s t test with Holm correction. Figure 2A – source data contains uncropped western blot images, Figure 2C-D – source data contains uncropped images of stained plates.

In parallel loss of function studies, Maf1 expression was decreased in ST2 cells using two different shRNAs (Fig. 3A). As expected, this resulted in an increase in pre-tRNA expression, particularly on day 10 (Fig. 3B). *Maf1* knockdown resulted in a decrease in Alp staining (Fig. 3C) and a robust reduction in mineralization (Fig. 3D). Osteoblast markers, namely *Sp7, Alp* and *Bsp* were significantly downregulated in cells (Fig. 3E).

**Figure 3.**
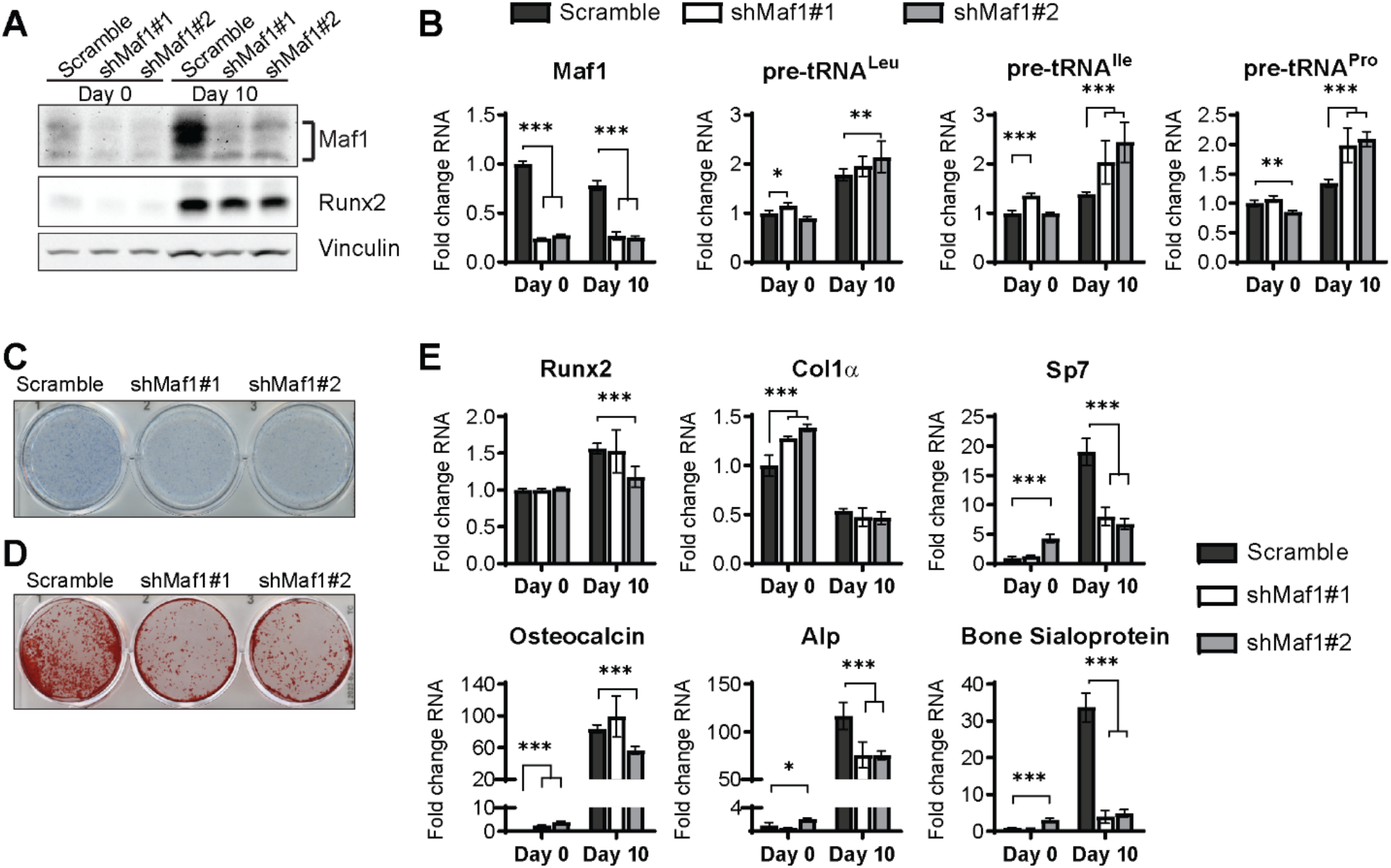
Maf1 knockdown decreases osteoblast differentiation of ST2 cells. (**A**) Western blot analysis showing Maf1, Runx2 and Vinculin expression in cells infected with a Scramble construct or Maf1 shRNA before, or 10 days after adding osteoblast differentiation medium. (**B**) qRT-PCR analysis of Maf1 and pre-tRNAs of ST2 cells expressing Scramble of shMaf1 before and on day after adding osteoblast differentiation medium. (**C**) Alkaline phosphatase staining of ST2 cells expressing scramble or lentiviral Maf1 shRNA after culture in osteoblast differentiation medium. (**D**) Alizarin red analysis of cells with scramble or Maf1 shRNA after culture in osteoblast differentiation medium. (**E**) qRT-PCR analysis showing relative expression of Runx2, Col1α, Sp7, Alp and Bone sialoprotein before, and 10 days after addition of osteoblast differentiation medium. Results represent means ±SD of three independent replicates, *P<0.05, **P<0.01, ***P<0.001 determined by Student’s t test with Holm correction. Figure 3A – source data contains uncropped western blot images, Figure 3C-D – source data contains uncropped images of stained plates.

Collectively, these results indicate that Maf1 functions to promote osteoblast differentiation. Reduced expression of Maf1 in ST2 cells and bone marrow stromal cells derived from *Maf1*^-/-^ mice resulted in a decrease in differentiation. Maf1 overexpression specifically in the mesenchymal cells of the long bones, increased bone mass. The paradoxical phenotype in *Maf1*^*-/-*^ mice therefore likely resulted from yet uncharacterized, non-cell-autonomous confounding effects on osteoblasts arising from global *Maf1* deletion.

Finally, Maf1 has been shown to promote adipogenesis since knockdown of *Maf1* in pre-adipocytes reduces adipocyte formation (29). We therefore examined how alterations in Maf1 expression may affect the differentiation of ST2 cells into adipocytes. We found that Maf1 overexpression produced an increase in oil red O stained cells and upregulated the expression of adipogenesis genes *Pparg, Cebpa* and *Fabp4* (Suppl. Fig 3). To determine if adipocyte formation was affected *Maf1* deficient mice, we isolated primary cells from *Maf1*^*-/-*^ mice. These cells showed increased tRNA transcription as expected and showed decreased differentiation into adipocytes as seen by Oil Red O stain (Suppl. Fig. 4A,B). Consistent with these results, histological analysis of femurs from 12-week-old *Maf1*^-/-^ mice showed that both adipocyte number and adipocyte volume was reduced (Suppl. Fig. 4C). These results confirm that in addition to promoting osteogenesis, Maf1 enhances adipocyte differentiation.

### Maf1-independent approaches to repress RNA pol III-dependent transcription decrease osteoblast differentiation

As Maf1 functions as a repressor of RNA pol III-dependent transcription (13, 41), we determined if other approaches that inhibit RNA pol III-dependent transcription would produce a similar increase in osteoblast differentiation. ST2 cells were treated with ML-60218, a chemical inhibitor of RNA pol III (42) or with DMSO vehicle as the control. Cells were treated for three days, starting one day before the addition of differentiation media. Two days after the initiation of differentiation, ML-60218 was removed, and cells were allowed to differentiate without further manipulation. tRNA gene transcription was significantly reduced by ML-60218 treatment (Fig. 4A). However, in contrast to what was observed with Maf1 overexpression, reduction of RNA pol III transcription by ML-60218 resulted in a decrease in Alp and Alizarin red staining (Fig. 4B,C) and a significant reduction in the expression of osteoblast marker genes (Fig. 4D). This effect was not specific to ST2 cells, as the differentiation of primary stromal cells from C57/BL6 mice was also significantly reduced with ML-60218 treatment (Suppl. Fig. 5).

**Figure 4.**
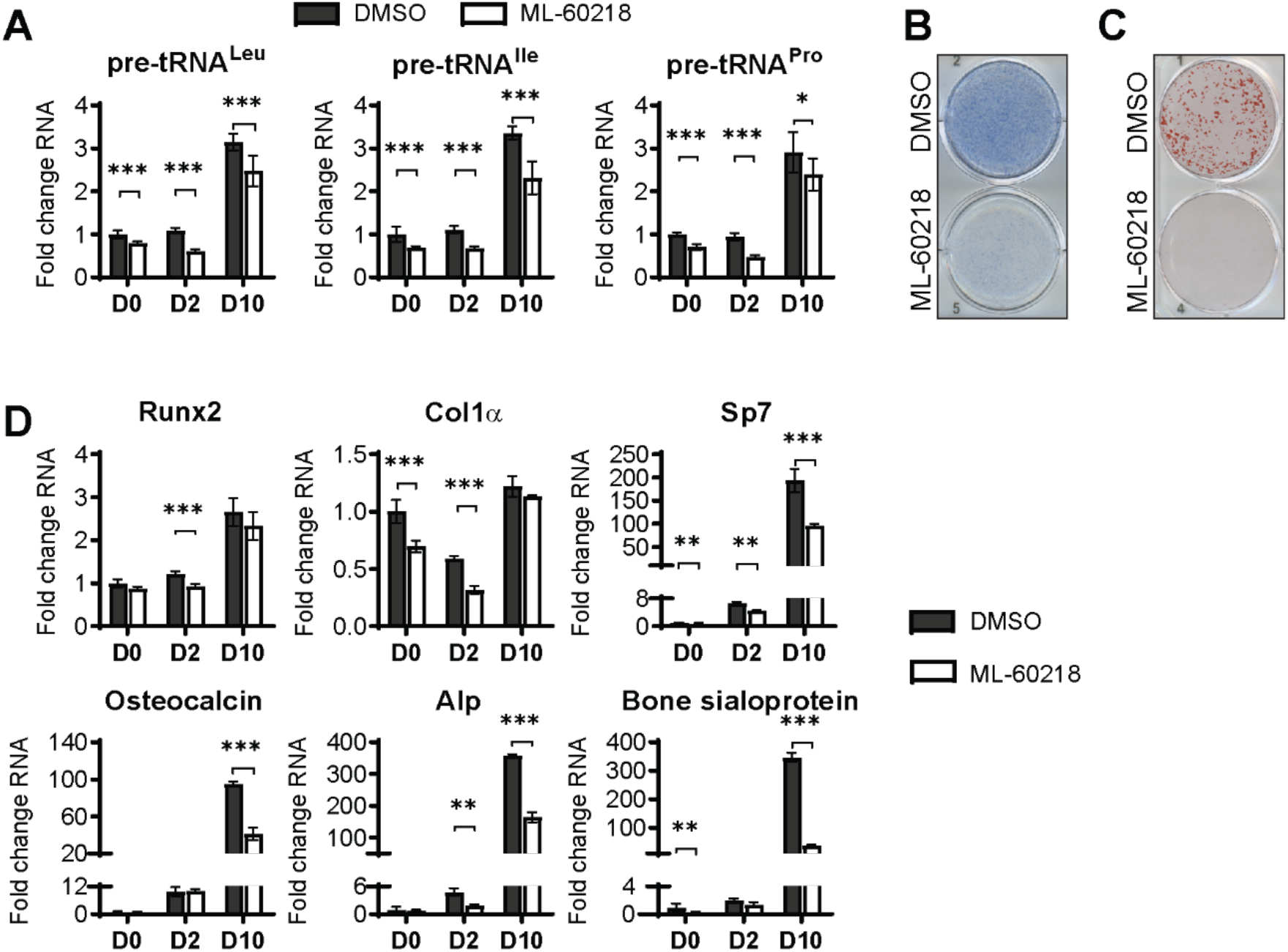
inhibition of RNA pol III-dependent transcription by ML-60218 decreases osteoblast differentiation and mineralization. ST2 cells were treated with 40µM ML-60218 for 3 days, starting at day -1 and differentiated into osteoblasts by addition of osteoblast differentiation medium on day 0. (**A**) qRT-PCR analysis of pre-tRNAs before and during differentiation after ML-60218 or DMSO treatment of ST2 cells. (**B**) Representative image of Alp staining of ST2 cells after osteoblast differentiation in DMSO or ML60218 treated cells. (**C**) Representative image of alizarin red analysis of ST2 cells after osteoblast differentiation and ML-60218 or DMSO treatment. (**D**) qRT-PCR analysis of Runx2, Col1α, Sp7, Osteocalcin, Alp and Bone Sialoprotein in ST2 cells on day 0, day 2 and day 10 during osteoblast differentiation. Results represent means ±SD of three independent replicates, *P<0.05, **P<0.01, ***P<0.001 determined by Student’s t test with Holm correction. Figure 4B-C – source data contains uncropped images of stained plates.

Using a complementary approach, we downregulated the expression of the RNA pol III-specific transcription factor *Brf1* to reduce RNA pol III transcription (Fig. 5A,B). Similar to ML-60218 treatment, *Brf1* knockdown decreased Alp and Alizarin red staining, as well as osteoblast marker expression (Fig. 5C-E). These results indicate that, while different approaches to decrease RNA pol III-dependent transcription all affect osteoblast differentiation, increased Maf1 expression functions to promote differentiation, while the other approaches used repress differentiation.

**Figure 5.**
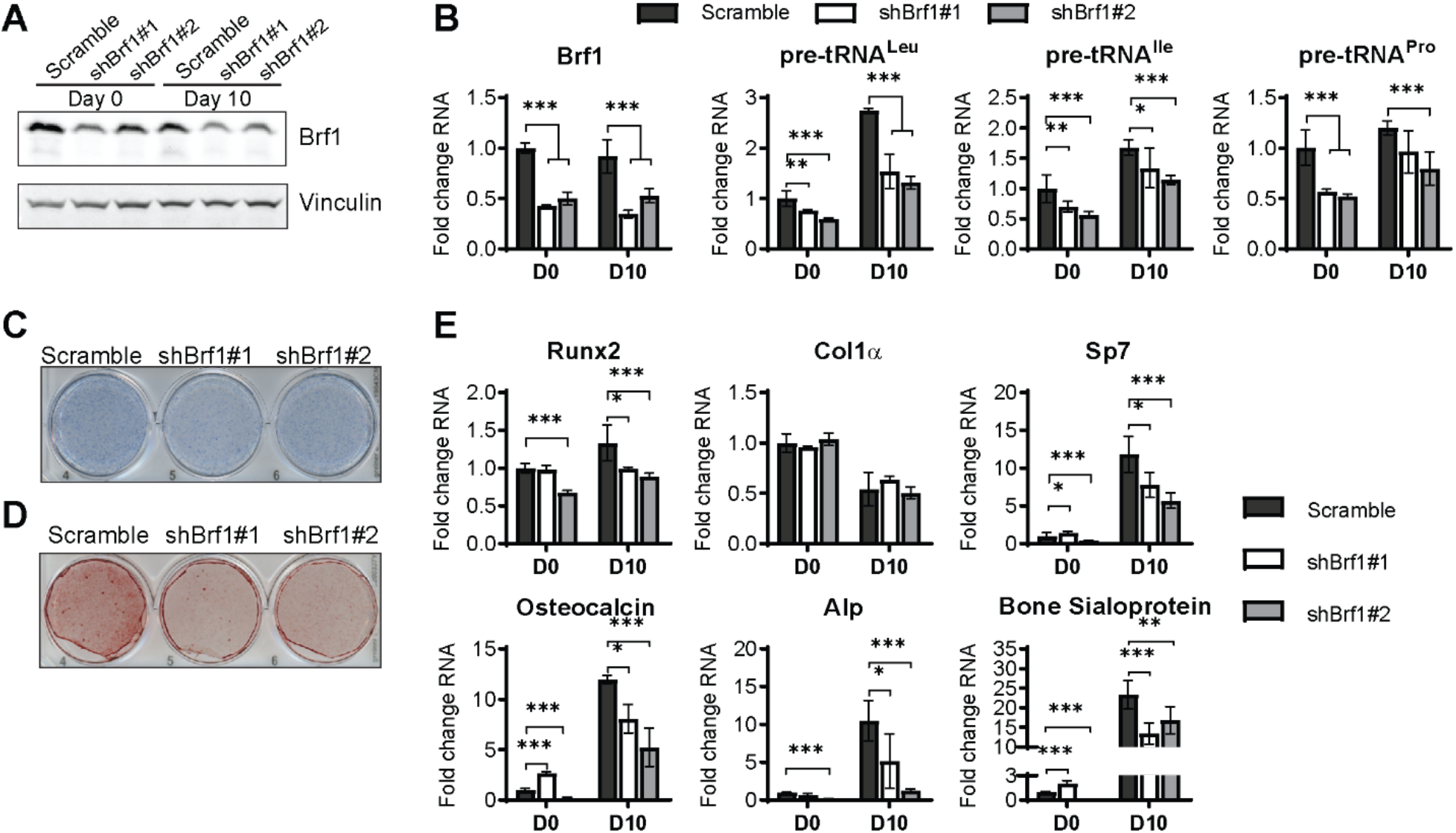
inhibition of RNA pol III-dependent transcription Brf1 knockdown decreases osteoblast differentiation and mineralization. ST2 cells were stably infected with scramble or Brf1 shRNA lentivirus and differentiated into osteoblasts by addition of osteoblast differentiation medium on day 0. (**A**) Western blot analysis showing Maf1, Runx2 and Vinculin expression in cells infected with a scramble construct or Maf1 shRNA before, or 10 days after adding osteoblast differentiation medium. (**B**) qRT-PCR analysis of Maf1 and pre-tRNAs of ST2 cells expressing Scramble of shMaf1 before and on day after adding osteoblast differentiation medium. (**C**) representative image of alkaline phosphatase staining of ST2 cells expressing scramble or lentiviral Maf1 shRNA after culture in osteoblast differentiation medium. (**D**) Representative image of alizarin red analysis of cells with scramble or Maf1 shRNA after culture in osteoblast differentiation medium. (**E**) qRT-PCR analysis showing relative expression of Runx2, Col1α, Sp7 (Osterix), Alp and Bone sialoprotein before, and 10 days after addition of osteoblast differentiation medium. Results represent means ±SD of three independent replicates, *P<0.05, **P<0.01, ***P<0.001 determined by Student’s t test with Holm correction. Figure 5A – source data contains uncropped western blot images, Figure 5C-D – source data contains uncropped images of stained plates.

To further delve into the opposing effects on osteoblast differentiation of Maf1 overexpression versus Brf1 downregulation or ML-60218, we examined the relative effects of the three perturbations on adipocyte differentiation from ST2 cells. Maf1-independent approaches to repress RNA pol III-dependent transcription produced an increase in adipogenesis observed upon oil red O staining and adipocyte marker expression (Suppl. Fig. 6,7). Interestingly, this was similar to what was observed by Maf1 overexpression. Thus, while all three repressors of RNA pol III transcription, ML-60218 treatment, *Brf1* knockdown and *Maf1* overexpression, enhance adipocyte differentiation, only Maf1 overexpression enhances osteoblast differentiation.

### RNA sequencing shows that different perturbations to alter RNA pol III transcription result in distinct gene expression profiles

To investigate the contrasting effect of altering Maf1 expression *versus* ML-60218 treatment or Brf1 knockdown on osteoblast differentiation, we performed RNA-seq on ST2 cells harvested before the start of differentiation, day 0, and at day 4. Each manipulation was compared to their own appropriate controls and triplicates were analyzed for each datapoint. Genes with an adjusted *P* value smaller than 0.05 and a log2 fold change greater than 0.7 in either direction were considered. We first compared changes in gene expression during osteoblast differentiation, without manipulating RNA pol III-dependent transcription. We examined potential changes in codon usage during osteoblast differentiation by comparing codon usage of upregulated genes, with an exhaustive list of gene coding sequences in mice. This revealed a significant bias towards the use of certain codons during differentiation (Fig. 6). Codon usage in our dataset was predominantly overlapping to codon bias in genes belonging to the GO term for osteoblast differentiation (GO:0001649). This suggests, for the first time, that codon bias may play a role during osteoblast differentiation.

**Figure 6.**
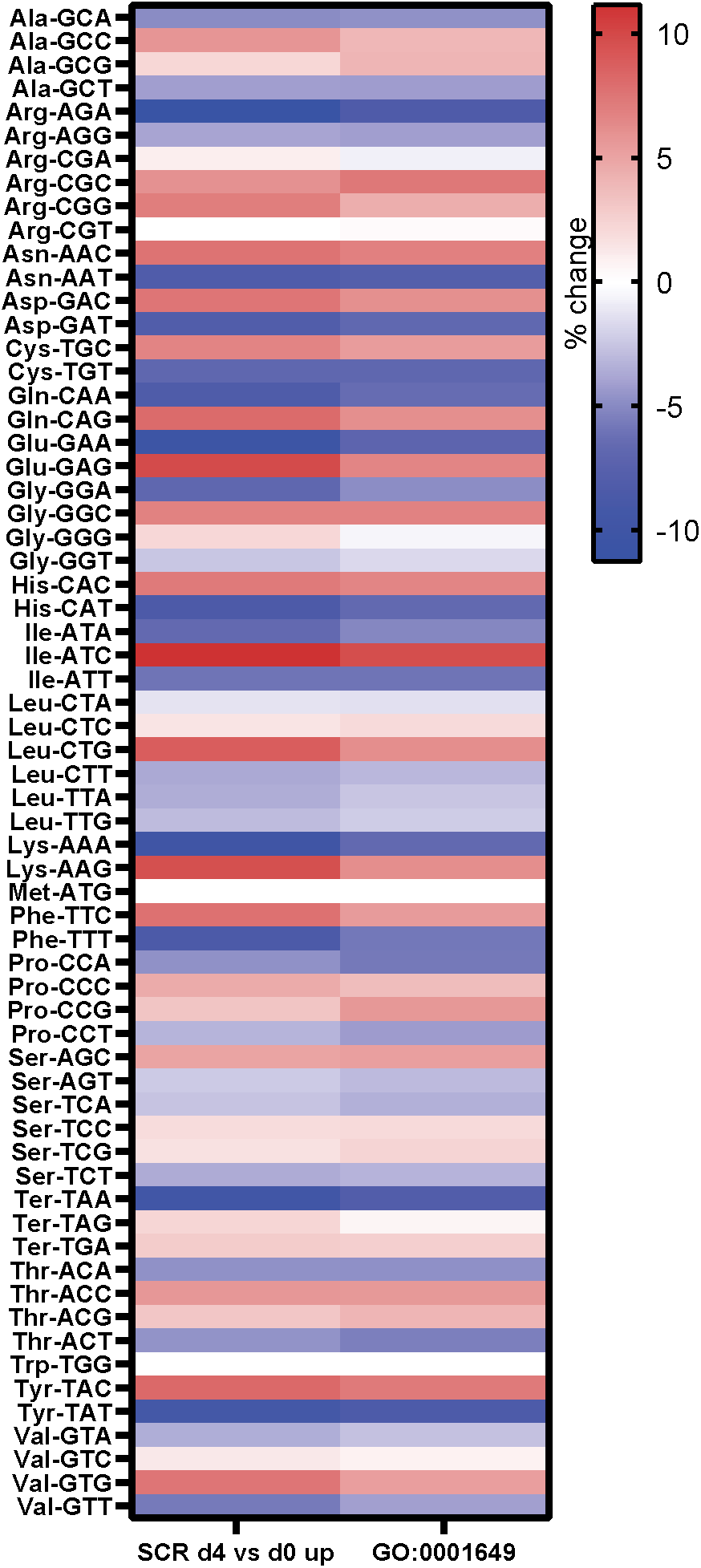
Genes expressed during osteoblast differentiation display significant codon bias. Relative changes in codon usage during osteoblast differentiation day 4, compared to day 0 for SCR control cells (left) or of genes that are members of the GO term 0001649 (osteoblast differentiation). Figure 6 – source data contains excel files with all codon analysis.

The interventions used to manipulate of RNA pol III-dependent transcription further resulted in distinct changes in gene expression on both day 0 and day 4 (Fig. 7A-B). To explore whether these changes relate to specific biological processes, we performed Gene ontology (GO) enrichment analysis on the data at day 0. Comparing the 20 most significantly enriched subgroups, we found that *Maf1* overexpression and *Brf1* knockdown resulted in enrichment for pathways expected to affect osteoblast differentiation, such as extracellular matrix organization and ossification. In contrast, ML-60218 treatment enriched mostly for lipid metabolism and adipocyte differentiation genes. The enrichment observed for these pathways without inducing differentiation suggests that manipulating RNA pol III-dependent transcription positions cells in a manner that affects their lineage determination. Our GO analysis also uncovered other biological processes that were changed although these and varied between the subgroups (Suppl. Fig. 8-11). Overall, these results reveal that different approaches to manipulate RNA pol III can produce disparate gene expression changes that lead to different biological outcomes.

**Figure 7.**
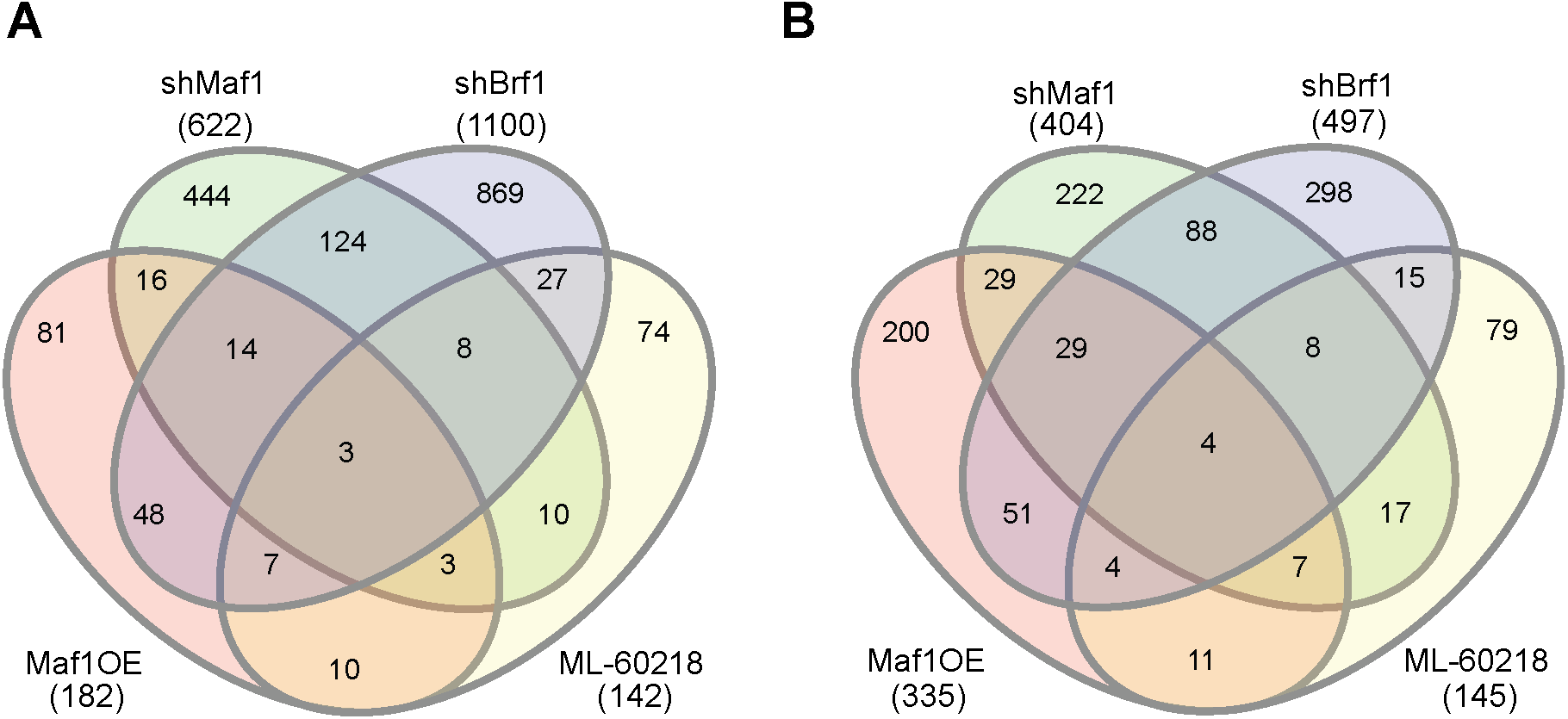
Manipulating RNA pol III in different manners results in distinct gene pools. Changes in gene expression were determined by Padj <0.05 and Foldchange > |log2 0.7|. Venn diagram showing overlap in gene changes on Day 0 (**A**) or (**B**) Day 4 (**B**). MAF1OE genes changes between pInd20-MAF1 and Pind20-Control; shMaf1 was compared to scramble control, shBrf1 was compared to scramble control; ML-60218 was compared to DMSO control. Figure 7 – source data contains excel files with all differentially expressed genes.

To determine which genes were specifically altered by *Maf1*, we compared changes in gene expression on day 0 by Maf1 overexpression and Maf1 knockdown. We only considered genes that were at least log2fold 0.7 changed in an opposing direction in each treatment set (Suppl. Fig. 12). This uncovered several genes that were not changed in the same direction by *Brf1* knockdown or ML-60218 treatment which have known effects on bone. Among these were phosphate regulating endopeptidase homolog, X linked (Phex), which increased upon *Maf1* overexpression and decreased by *Brf1* knockdown or ML-60218 treatment. Of note, *Phex* plays a key role in promoting bone mineralization and phosphate homeostasis (43). In addition, *Col15a1*, which is associated with early osteoblast differentiation (44), and *Lysyl oxidase like 2* (*Loxl2*), which is involved in collagen crosslinking (45), were also increased by *Maf1* overexpression. In contrast, *Rhomboid 5 homolog 2* (*Rhbdf2*) expression was decreased. *Rhbdf2* knock out mice display a high bone mass phenotype (46) suggesting RHBDF2 may play a role in regulating bone mass. Together, the results suggest that Maf1 may specifically regulate a subset of genes that play a role in regulating bone mass.

## Discussion

Maf1 is a key repressor of transcription by RNA pol III (13, 14). Changes in Maf1 have been shown to enhance adipocyte differentiation (29)(Suppl. Fig. 5 and 6). *Maf1*^*-/-*^ mice have a short and lean phenotype with resistance to diet-induced obesity, decreased fertility and fecundity, and increased longevity/healthspan (21). Here, we demonstrate that these mice display an increase in bone volume. However, *Maf1*^*-/-*^ derived primary stromal cells showed a decrease in osteoblast formation in *ex vivo* cultures. This latter finding was consistent with the effects of Maf1 overexpression specifically in the mesenchyme of long bones, which resulted in enhanced osteoblast differentiation and an increase in bone mass. In these latter mice, compared with *Maf1*^*-/-*^ mice, any confounding actions due to the absence of Maf1 in other cells and tissues, such as possible endocrine or paracrine effects, are limited. Thus, our results indicate that the ability of Maf1 to regulate bone mass involves both cell autonomous and non-cell-autonomous actions. This idea is further corroborated by our *in vitro* results showing that osteoblast differentiation is regulated by increasing or decreasing *Maf1* expression in ST2 cells, suggesting that Maf1 promotes osteoblast differentiation and mineralization. As Maf1 also enhances adipogenesis(29)(Suppl. Fig. 1), we establish Maf1 as an important regulator in the development and differentiation of mesenchymal cells into multiple lineages.

Maf1 is a well-established repressor of RNA pol III-mediated transcription through its direct interaction with RNA pol III (14, 15). To further determine whether Maf1 functions to promote osteoblast differentiation through its ability to regulate RNA pol III-dependent transcription, we used complementary approaches to repress this transcription process. During differentiation, we observed an overall increase in the tRNA gene transcripts, which were analyzed. This increase was repressed by *Maf1* overexpression, *Brf1* downregulation, or chemical inhibition of RNA pol III. Surprisingly, however, in contrast to the positive regulation of osteoblast differentiation by Maf1, chemical inhibition of RNA pol III or *Brf1* knockdown resulted in a decrease in osteoblast differentiation. Thus, while different perturbations in RNA pol III-dependent transcription all affect the differentiation process, Maf1-mediated changes produce an opposing action compared with chemical inhibition of RNA pol III or decreased *Brf1* expression. This is in contrast to what we observed for adipogenesis, where all three approaches to repress RNA pol III transcription similarly increased adipocyte formation (Suppl. Fig. 3,4,6,7; (29).

To understand the basis of the different osteoblast differentiation outcomes, we examined changes in gene expression that resulted from the different perturbations of RNA pol III-dependent transcription. RNA seq revealed that alterations in RNA pol III-mediated transcription prior to, and during differentiation, result in a limited number of overlapping changes, with the four conditions largely producing distinct changes in gene expression profiles. However, changes in several established regulators of osteoblast formation and function were identified that correlated with the differentiation outcomes. Overall, the different gene expression profiles likely contribute to the differences in osteoblast differentiation that we observe. However, the precise mechanism underlying the difference between Maf1-regulated changes compared with the alternate approaches to repress RNA pol III-mediated transcription remains unclear. Unlike RNA pol III and Brf1, Maf1 is not an essential RNA pol III transcription factor and it functions to repress this transcription process by directly interacting with RNA pol III (14). Given that Maf1 is recruited to certain RNA pol II promoters, (13, 16, 17, 47), it is conceivable that the ability of Maf1 to regulate both RNA pol III- and RNA pol II-transcribed genes contributes to its ability to drive osteoblast differentiation. This may also explain its differential effect on osteoblast differentiation when compared with the alternative approaches used to selectively repress RNA pol III-dependent transcription.

RNA-seq analysis revealed distinct changes in gene expression exhibited by chemical inhibition of RNA pol III and *Brf1* knockdown, despite similar outcomes on osteoblast differentiation. These data indicate that manipulating RNA pol III transcription using different approaches results in disparate outcomes on gene expression. RNA pol III transcribes a variety of untranslated RNAs. While changes in any of these targets could potentially affect mRNA expression, this could be due to differential changes in the tRNA population. The prevalence of specific tRNAs has been correlated with codon biased translation in multiple tissues (48– 51) and our analysis of mRNA changes during osteoblast differentiation showed that a significant codon bias emerges during this process. Thus, it is conceivable that during osteoblast differentiation, changes in tRNAs are required to efficiently drive codon biased translation of mRNAs needed for differentiation to proceed. There are several other mechanisms that could contribute to the observed differences in gene regulation by RNA pol III-mediated transcription. Changes in other RNA pol III-derived transcripts, the generation of tRNA fragments (52, 53), or regulation of RNA pol II genes through the recruitment of RNA pol III to nearby SINE sites such as described for *CDKN1a* (19), could all potentially contribute to the differential expression of osteoblast genes when RNA pol III-mediated transcription is altered. Future work will be needed to identify the specific mechanisms by which Maf1 and RNA pol III-mediated transcription alter gene expression to regulate osteoblast differentiation.

In all, our results describe a novel role for Maf1 in bone biology. Given that different approaches used to modulate RNA pol III-dependent transcription also affect osteoblast differentiation, albeit in an opposing direction from Maf1-mediated effects, this supports the idea that Maf1 functions, at least in part, to regulate osteoblast development and bone mass through its ability to control RNA pol III-mediated transcription. Distinct qualitative or quantitative changes resulting from different perturbations in RNA pol III-dependent transcription may also play a role in developmental disorders. Interestingly, several different syndromes relating to mutations in RNA pol III subunits show very heterogeneous phenotypes. This includes *POLR3*-related hypomyelinating leukodystrophies (54–56), Wiedemann-Rautenstrauch syndrome, a neonatal progeroid syndrome (57–60) and Cerebellar hypoplasia with endosteal sclerosis (37, 38). Additionally, *BRF1* mutations have been shown to be causative for cerebellofaciadental syndrome (34, 35, 61). Some patients with RNA pol III-related mutations, but not all, show bone-related phenotypes (34–38). The considerable heterogeneity of these syndromes has been suggested to be related to differential changes in RNA pol III-dependent transcription (55). Thus, some cells and tissues, including bone and its cells, may be more sensitive to disruption of RNA pol III-mediated transcription. Together, these collective findings and our current study indicate that the exquisite regulation of RNA pol III plays an essential role in a variety of biological and developmental processes.

## Materials and methods

### Mouse lines and bone analyses

All mouse experiments were performed according to a protocol approved by the Institutional Animal Care and Use Committee at Baylor College of Medicine and Albert Einstein College of Medicine. *Rosa26*-Lox-stop-lox-Maf1-HA (LSL-Maf1) mice were generated by injecting an engineered construct into mouse (C57Bl6/J strain) embryonic stem cells and selecting for homologous recombinant clones. The selected clones were used to generate chimeric mice by blastocyst injection. The chimeric mice were bred to found the LSL-MAF1 colony. *Prx1*-Cre lines were a kind gift from Dr. Brendan Lee. LSL-MAF1 mice were mated with *Prx1*-Cre for conditional overexpression of MAF1-HA. Littermate controls expressing only Cre were used as a control. Left femurs were collected for µCT at 12 weeks. They were fixed for 48 hours in 4% paraformaldehyde (PFA) and stored at 4°C in 70% ethanol. µCT of left femurs was performed using the Scanco µCT-40 system at 16 µm resolution. 75 slices in the metaphyseal region of each femur was analyzed, starting at 10 slices beyond disappearance of the growth plate. Tibiae and right femurs were dissected, bone marrow was washed out and bones were subsequently snap frozen in liquid nitrogen for subsequent protein and RNA isolation. For the *Maf1*^-/-^ mice, µCT measurements at the spine, femurs and tibiae were performed through the courtesy of Dr. Jay Cao (USDA, North Dakota) using a Scanco µCT-40 scanner.

### ST2 cell culture and differentiation

ST2 cells were acquired from RIKEN BRC cell bank. Cells, which tested negative for mycoplasma, were grown in basic medium, ascorbic acid free α-MEM (Caisson laboratories) supplemented with 10% FBS (Gibco). For the knockdown experiments, cells were either infected with a scrambled control (gift from Sheila Stewart addgene #17920 (62), *Maf1* shRNA (#1 TRCN0000125776 and #2 TRCN0000125778) or *Brf1* shRNA (#1 TRCN0000119897 or #2 TRCN0000119901). pInd20-Maf1-HA was cloned by taking Maf1-HA from pFTREW-Maf1-HA (16) into a pInducer20 construct by gateway cloning using LR clonase (Thermo Fisher). The empty pInducer20 vector was a gift from Stephen Elledge (Addgene #44012) (63). Virus production and cell infection was performed as described previously (29). Cells were used for differentiation within 3 passages of selection. For Maf1 overexpression, pInducer20-MAF1HA infected, or pInducer20-empty cells were treated with 1 µM doxycycline 24 h before differentiation was started. For ML-60218 treatment, cells were treated starting 24 h before differentiation with 40 µM ML-60218 in DMSO (Millipore) or an equal volume of DMSO as control. ML-60218 treatment continued for 2 days after differentiation was initiated after which the compound was removed. For osteoblast differentiation, ST2 cells were plated at 1.8×10^5^ cells *per* well in a 6-well plate. Cells were grown to confluence after which osteoblast differentiation medium, basic media with 50µg/mL ascorbic acid and 10mM β-glycerolphosphate was added (day 0) and changed every two days. For adipocyte differentiation, ST2 cells were plated at 1.8×10^5^ cells *per* well in a 6-well plate and grown to confluency. On day 0, adipogenic medium was added (basic media with 1µM rosiglitazone, 0.5 mM 3-isobutyl-1-methyl xanthine, 2 µM dexamethasone and 10 µg/mL insulin). After 2 days, the media was changed to maintenance medium (basic media with 10 µg/mL insulin), which was changed every two days for the remainder of the experiment.

### Osteoclast Cultures

Bone marrow cells were isolated from femora and tibiae of *Maf1*^*-/-*^ and wild type mice in alpha-MEM. Cells were cultured for 2 days with M-CSF (30 ng/ml). Non-adherent cells were collected and purified by Ficoll-Plus (Amersham Pharmacia, Arlington Height, IL). They were then incubated with M-CSF (30 ng/ml) and RANK-L (100 ng/ml) for 4 to 6 days followed by staining for Tartrate-resistant acid phosphatase (TRAP) using a kit (Sigma) *per* manufacturer’s instruction. The number of TRAP-positive cells was counted.

### Cfu-f and Cfu-ob Cultures

Marrow stromal cells were cultured in the presence of ascorbate-2-phosphate (1 mM) (Sigma). Colony forming units-fibroblastoid (Cfu-f) and colony forming units-osteoblastoid (Cfu-ob) were counted, respectively, following alkaline phosphatase staining after 14-day cultures, or von Kossa staining after 21 day cultures.

### Primary stromal cell culture

Primary bone marrow stromal cells were isolated from femurs and tibiae of 6-8-week-old mice. Bones were dissected, cleaned and the marrow was flushed out. Bone pieces were digested using 2.5% collagenase IV (Gibco) for 2-4 hours at 37°C. Cells were strained and maintained in basic medium with 10ng/mL FGF2 (Biovision). For differentiation, cells were plated in a 48 or 6 well plates and differentiation was performed as described above.

### RNA isolation and quantitative PCR

Total RNA from cells was isolated using the quick-RNA miniprep kit (Zymo Research) following manufacturer’s protocol. For femurs, samples were ground using mortar and pestle in liquid nitrogen, and then further disrupted in RNA stat-60 (Tel-Test Inc.) using a polytron. RNA was isolated using the Direct-zol miniprep kit (Zymo Research). cDNA was synthesized using Superscript IV First Strand Synthesis Kit (Invitrogen). Quantitative PCR was performed using SYBR fast qPCR mastermix (KAPA Biosystems) on the Roche 480 Lightcycler. Gene specific primers are described in Suppl. Table 2. RNA was quantified relative to *Ef1a* for osteoblast differentiation and *Ppia1* for adipocyte differentiation.

### Protein isolation

Cells were washed twice and lysed in RIPA buffer. Tibia were ground in mortal and pestle, and further disrupted in RIPA buffer using a polytron. Samples were then sonicated. Cell lysate concentrations were measured using DC protein assay (Biorad) and similar amounts of protein lysate were loaded. The following antibodies were used: Maf1 (H2), TFIIIB90 (Brf1) (A8), Vinculin (7F9) and β-actin (C4) (Santa Cruz), Runx2, Pparγ, Fabp4 (Cell signaling) and HA (Roche).

### Staining

Cells were fixed for 10 minutes in 4% PFA, washed twice with PBS and once with water. Oil red O staining was performed using 0.3% Oil Red O solution (Sigma). Alkaline phosphatase staining used an alkaline phosphatase blue substrate kit (Vector Laboratories). Alizarin red staining was performed using 1% alizarin red (Sigma-Aldrich) at a pH of 4.2. Cell counts were taken using the Cytation 5 Microscope. Alizarin Red was extracted using 10% cetylpyridinium chloride (CPC) and absorption was measured at 563 nM.

### Sequencing

ST2 cells were prepared, plated, and differentiated as described above. shBrf1#2 and shMaf1#2 were used for sequencing analysis. Triplicates of each sample were used and RNA was extracted on day 0 and day 4. For each replicate, three wells of a 6-well plate were combined. For each condition triplicate RNA was isolated using the quick RNA miniprep kit (Zymo Research). Library preparation and RNA-seq was performed by by Novogene Co. (Sacramento CA, USA). Differentially expressed genes were determined using DESeq2 with FDR <0.05 and |log2foldchange| > 0.7. One replicate of sh*Brf1* at day 0 was considered an outlier by principal component analysis and hierarchical clustering and removed from analysis. (GO) analysis was performed using the clusterProfiler R package (64). Venn diagrams were made using interactivenn (65). For codon usage analysis an exhaustive list of gene coding sequences was obtained from GENCODE (M27) and codon use rates were calculated. For each codon, a selection rate against other potential isodecoders was determined. Gene subsets were established from alteration in RNAseq at over log2foldchange 0.7 in either direction and padj < 0.05 by DESeq2, or by membership in gene ontology as Osteoblast Differentiation. For comparisons between two groups two tailed students t-test were performed followed by Benjamini-Hochberg correction of the full comparison set.

### Statistical analysis

For comparisons between two groups two tailed students t-test were performed. For comparisons with more than two groups, ANOVA was used followed by paired t-tests with Holm correction. Significance was determined at *P*<0.05.

## Acknowledgments

We would like to than the Genetically Engineered Rodent Models Core at the Baylor College of Medicine for assistance with mouse model production. Resources accessed through the core were supported by a National Institutes of Health grant (P30CA125123) to the Dan L. Duncan Comprehensive Cancer Center. We would like to thank Brian Dawson at Baylor College of Medicine for his help with the µCT measurements. This work was supported by NIH grants R01 CA108614 and R01 CA74138 (to D.L.J), R01 GM120358 (to I.M.W.), NIH--U19 AG060917 (M.Z.), R01 AG071870, U01 AG073148 and R01AG074092 (to M.Z. and T.Y.). M.Z. also thanks the Harrington Discovery Institute for the Innovator–Scholar Award.

## Figures

**Supplemental Figure 1.**
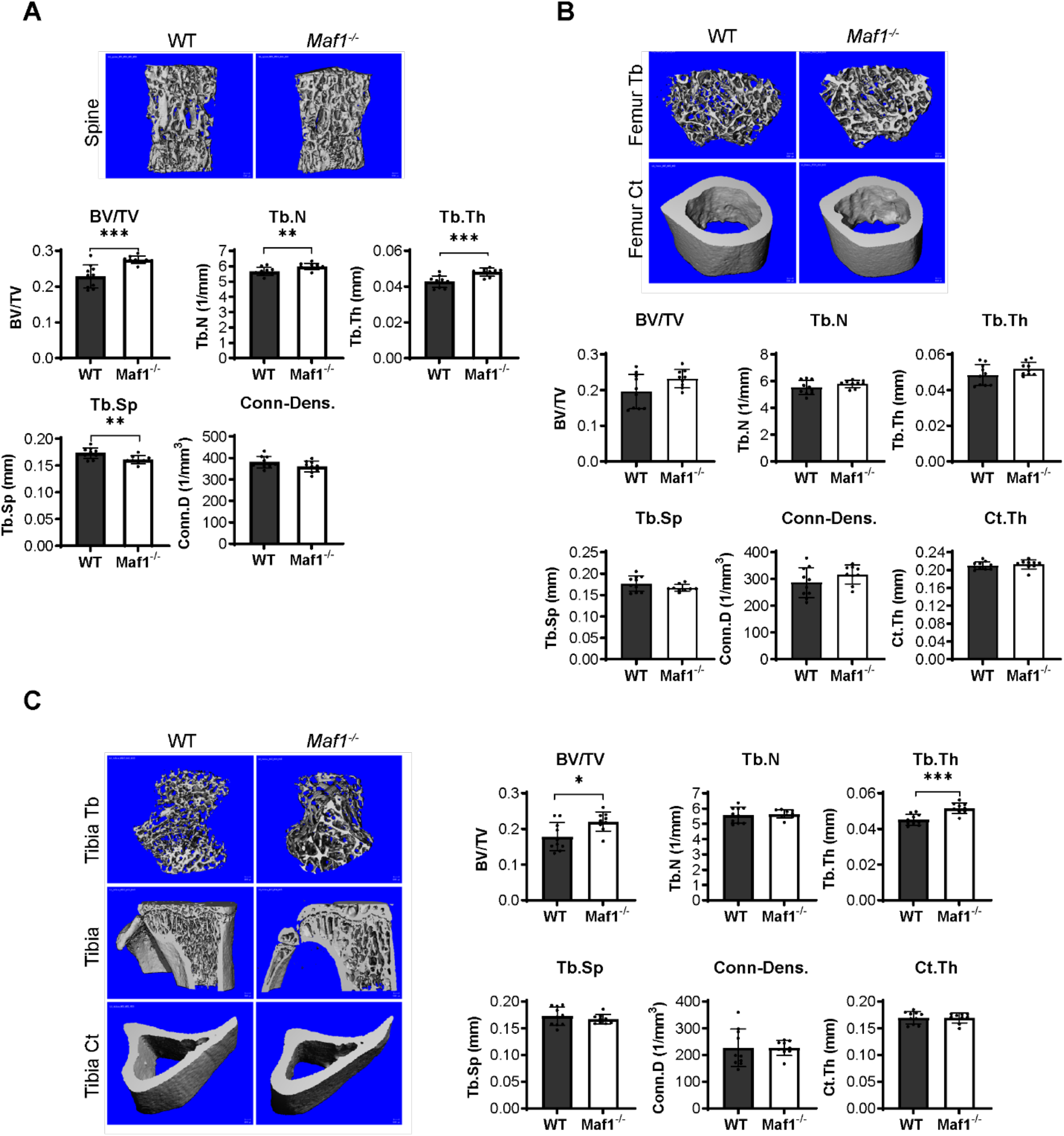
*Maf1*^*-/-*^ mice show increased bone mass in the spine. Spines and femurs were taken from 12-week-old male *Maf1*^*-/-*^ mice or their WT counterparts. µCT measurements from the spine (**A**) the femoral bone (**B**), or the tibia (**C**). Representative images of µCT of the spine (top) (**A**), femur (top) (**B**), or tibia (left) (**C**). Quantification of µCT analysis (bottom for Spine (**A**), femur (**B**) or right (**C**): Bone volume/total volume (BV/TV), trabecular number (Tb.N), trabecular thickness (Tb.Th), trabecular separation (Tb.Sp), connectivity density (Conn-Dens.) and cortical thickness (Ct.Th). WT n=10 *Maf1*^*-/-*^ n=9. Results represent means ±SD, *P<0.05, **P<0.01, ***P<0.001 determined by Student’s t test.

**Supplemental Figure 2.**
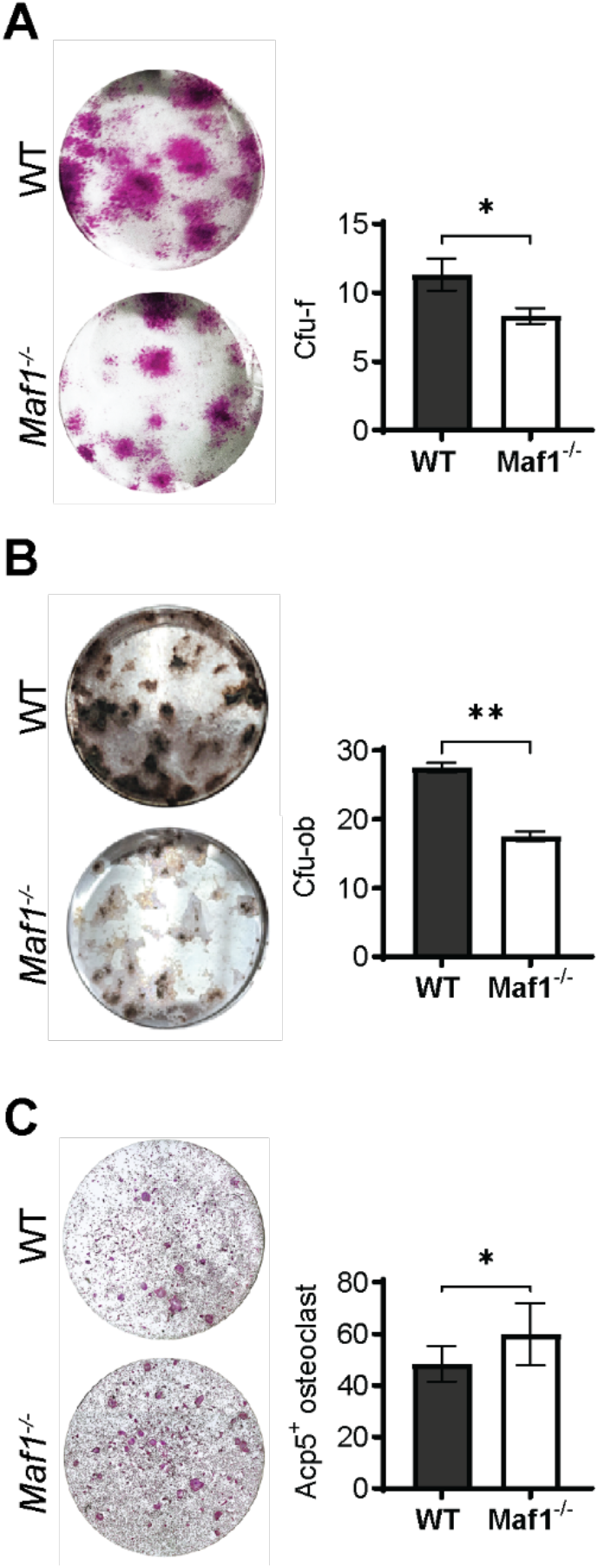
*Ex vivo* analysis *Maf1*^*-/-*^ cells show increased osteoblast differentiation and decreased osteoclast formation. (**A**) Representative image (left) and quantification (right) of alkaline phosphatase labeled colony forming units-fibroblastoids (Cfu-F), right quantification. (**B**) Representative image (left) and quantification (right) of Von Kossa labeled colony forming units-osteoblastoid (Cfu-ob). (**C**) Representative image (left) and quantification (right) of Acp5^+^ cells after osteoclast differentiation using 100ng/mL Rank-l. Results represent means ±SD, *P<0.05, **P<0.01 determined by Student’s t test.

**Supplemental Figure 3.**
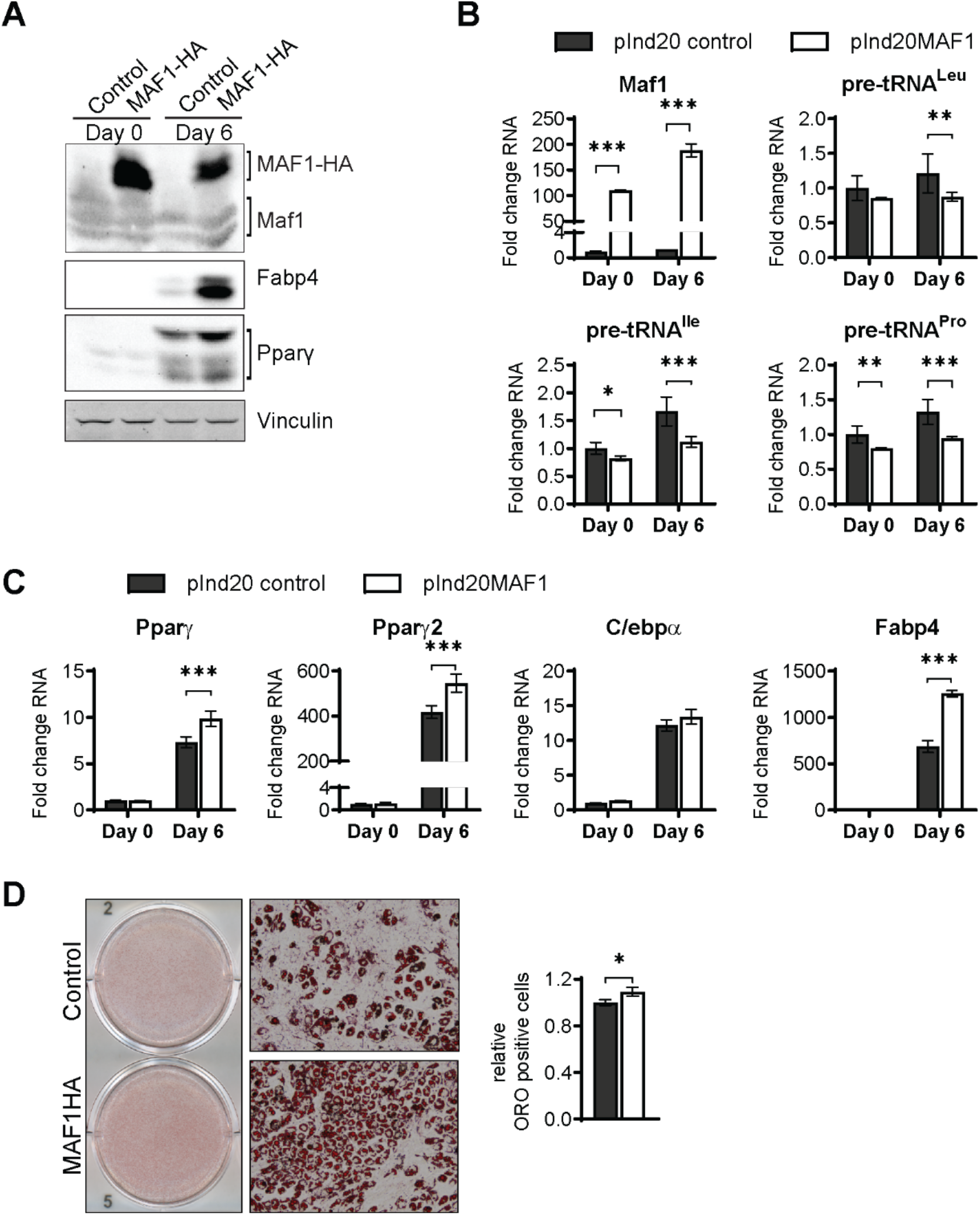
MAF1 overexpression enhances adipogenesis in ST2 cells. MAF1 or a control vector were expressed in ST2 cells and cells were subsequently differentiated into adipocytes as described in material and methods. (**A**)Western blot analysis shows MAF1, Fabp4, PPARγ and vinculin expression on day 0 and day 6 of adipocyte differentiation. (**B**) qRT-PCR analysis of MAF1 and pre-tRNAs in control and MAF1 overexpressing ST2 cells before and during adipocyte differentiation. (**C**) qRT-PCR of Pparγ, Pparγ2, C/ebpα and Fabp4 of ST2 cells expressing a control or MAF1-HA vector before and after adipocyte differentiation. (**D**) Representative images of Oil red O staining of adipocytes differentiated from ST2 cells expressing control or MAF1-HA (left), 10X images (middle) quantification of Oil red O positive cells (right). Results represent means ±SD of three independent replicates, *P<0.05, **P<0.01, ***P<0.001 determined by Student’s t test with Holm correction. Supplementary Figure 3A – source data contains uncropped images of western blot analysis. Supplementary Figure 3D – source data contains uncropped images of the Oil red O-stained cells, additional 10x images and stitched images at 4x used for analysis.

**Supplemental Figure 4.**
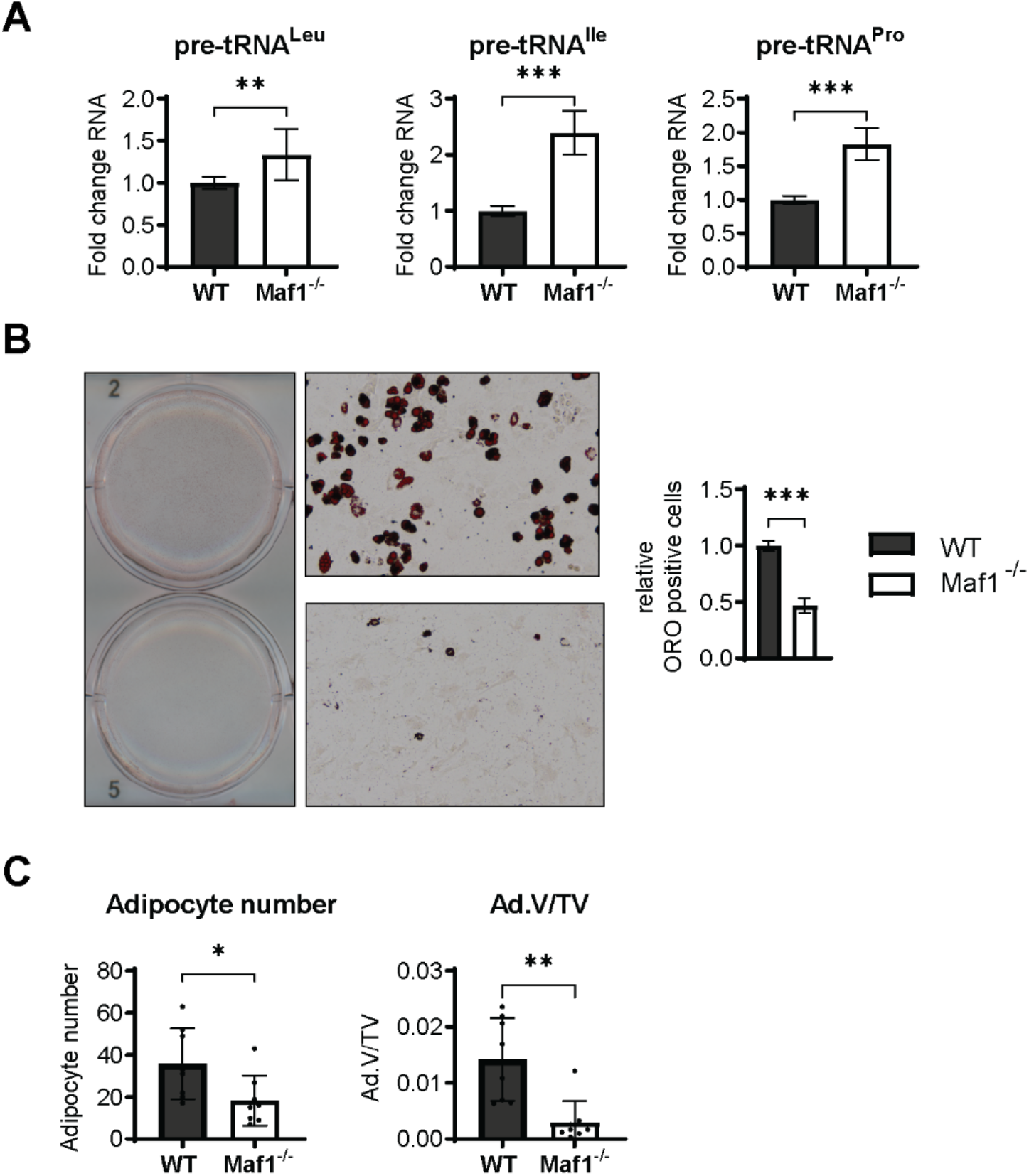
Maf1 deficiency decreases adipocyte differentiation *in vitro* and bone marrow adipocytes *in vivo*. Primary stromal cells isolated from femurs of 6–8-week-old Maf1^-/-^ or WT male mice. (**A**) qRT-PCR analysis of pre-tRNAs in WT or *Maf1*^*-/-*^ cells. Results from 12 independent replicates (**B**) Oil Red O staining of WT and *Maf1*^*-/-*^ cells differentiated into adipocytes for 9 days. Representative image (left), 10X images (middle) quantification of Oil red O positive cells (right). Results of 3 independent replicates. (**C**) Histological analysis of 12-week-old femurs of WT and *Maf1*^*-/-*^ mice. Adipocyte number and adipocyte volume/ total volume (Ad.V/TV). n=8 for WT and n=8 for *Maf1*^*-/-*^ mice femurs. Results represent means ±SD, *P<0.05, **P<0.01, ***P<0.001 determined by Student’s t test. Supplementary Figure 4B – source data contains uncropped images of the Oil red O-stained cells, additional 10x images, and stitched images at 4x used for analysis.

**Supplemental Figure 5.**
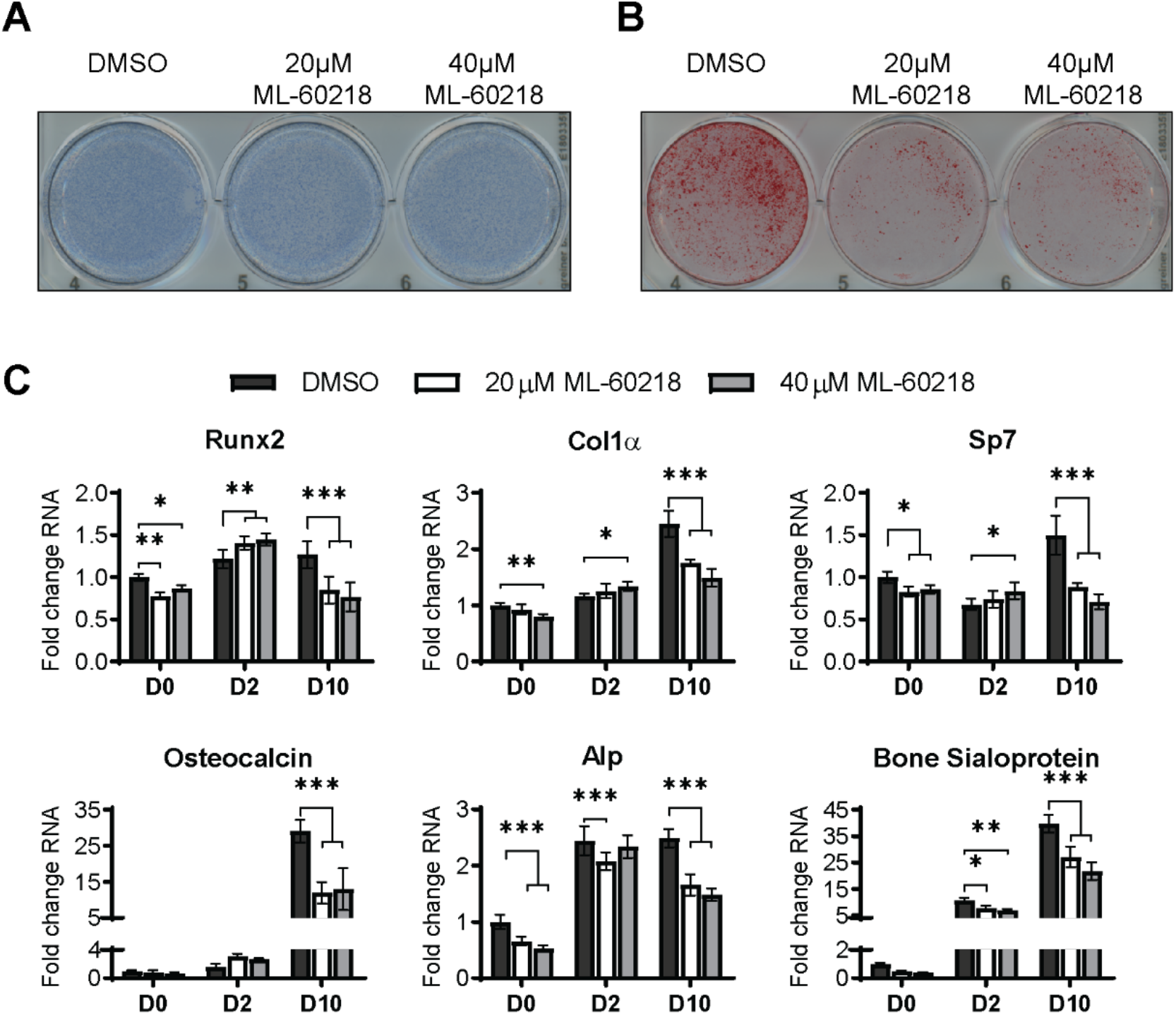
ML-60216 treatment decreases osteoblast differentiation of primary stromal cells. Primary stromal cells isolated from 6–8-week-old C57/BL6 WT mice were treated with ML-60218 for 3 days before, and during differentiation into osteoblasts by addition of osteoblast differentiation medium on day 0. (**A**) Representative image of Alp staining of ST2 cells after osteoblast differentiation of DMSO or ML60218 treated cells. (**B**) Representative image of alizarin red analysis of ST2 cells after osteoblast differentiation and ML-60218 or DMSO treatment. (**C**) qRT-PCR analysis of Runx2, Col1α, Sp7, Osteocalcin, Alp and bone sialoprotein expression relative to β-actin in primary stromal cells on day 0, day 2 and day 10 during osteoblast differentiation. Results represent means ±SD of three independent replicates, *P<0.05, **P<0.01, ***P<0.001 determined by Student’s t test with Holm correction. Supplementary Figure 5A-B – source data contains uncropped images of stained plates.

**Supplemental Figure 6.**
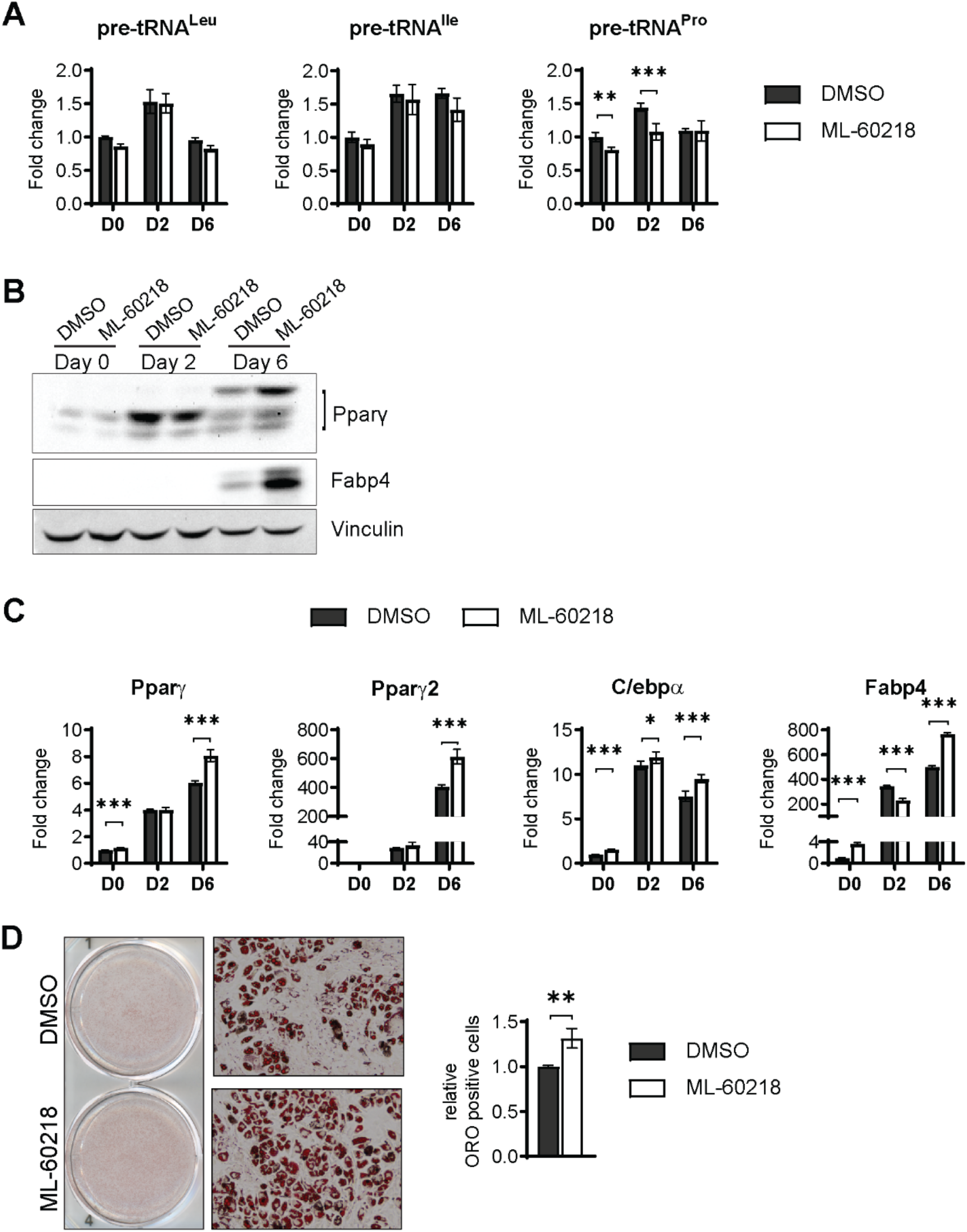
ML-60218 treatment enhances adipogenesis of ST2 cells. ST2 cells were treated for 3 days with 40µM ML-60218 or DMSO between day -1 and day 2 of adipocyte differentiation. (**A**) qRT-PCR analysis of pre-tRNA expression before and during adipocyte differentiation. (**B**) Western blot analysis of Pparγ, Fabp4 and Vinculin. (**C**) qRT-PCR analysis of adipocyte markers Pparγ, Pparγ2, C/ebpα and Fabp4. (**D**) Oil red O staining of adipocytes on day 8 of adipocyte differentiation. Representative wells (left), representative 10X microscope image (middle), relative oil red O positive cells as determined by citation 5 scanning of 3 wells (right). *P<0.05, **P<0.01, ***P<0.001 determined by Student’s t test with Holm correction. Supplementary Figure 6B – source data contains uncropped images of western blot analysis. Supplementary Figure 6D – source data contains uncropped images of the Oil red O-stained cells, additional 10x images and stitched images at 4x used for analysis.

**Supplemental Figure 7.**
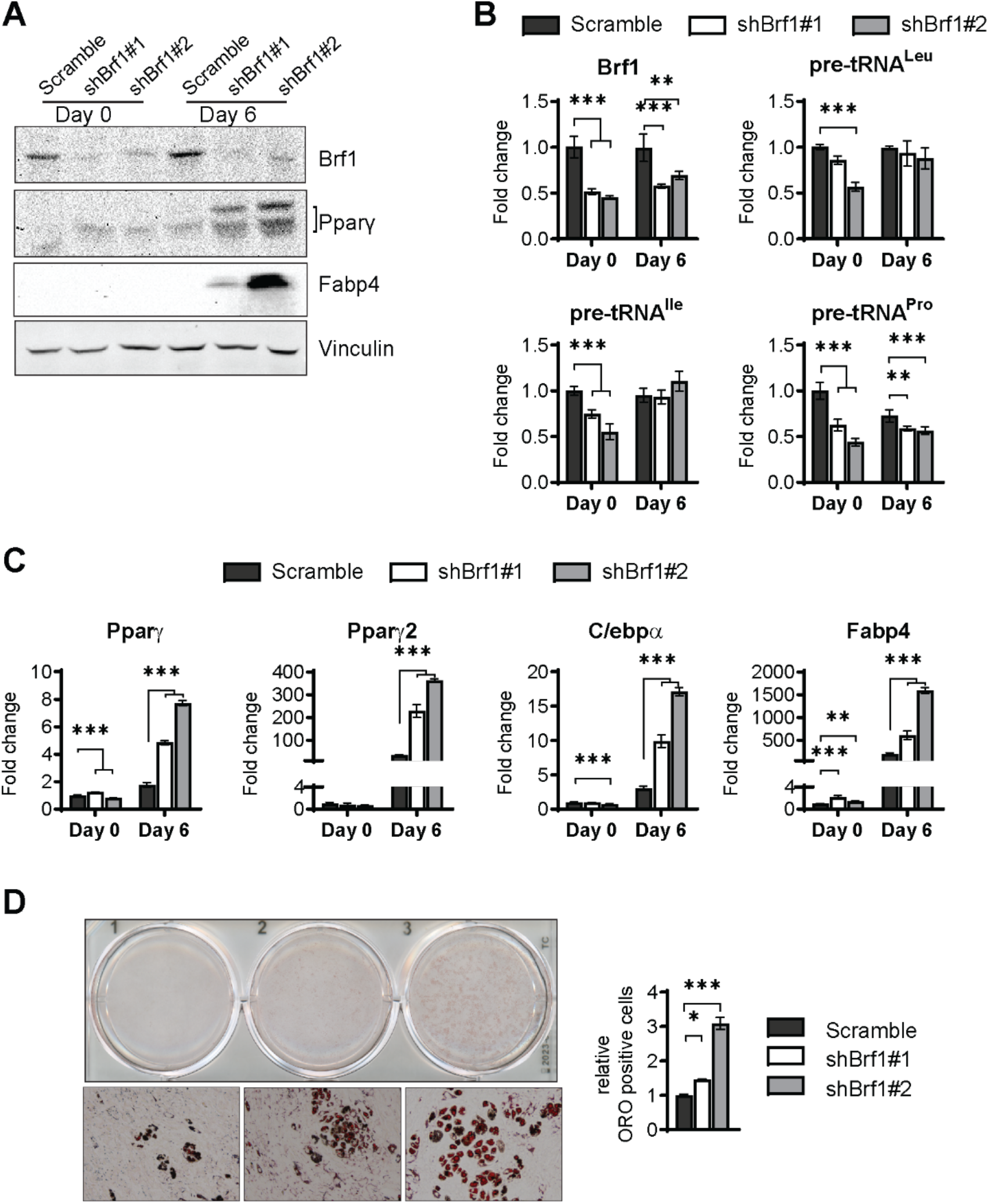
Brf1 knockdown enhances adipogenesis in ST2 cells. ST2 cells were stably infected with scramble or two different shBrf1 constructs and differentiated into adipocytes as described in “Material and Methods”. (**A**) Western blot analysis of Brf1, Pparγ, Fabp4 and vinculin expression on day 0 and day 6 of adipocyte differentiation. (**B**) qRT-PCR analysis of Brf1, and pre-tRNA expression during adipocyte differentiation. (**C**) qRT-PCR analysis of adipocyte markers Pparγ, Pparγ2, C/ebpα and Fabp4. (**D**) Oil red O staining of adipocytes on day 8 of adipocyte differentiation. Representative wells (top), representative10X microscope image (bottom), relative oil red O positive cells as determined by citation 5 scanning of 2 wells (right). *P<0.05, **P<0.01, ***P<0.001 determined by Student’s t test with Holm correction. Supplementary Figure 7A – source data contains uncropped images of western blot analysis. Supplementary Figure 7D – source data contains uncropped images of the Oil red O-stained cells, additional 10x images and stitched images at 4x used for analysis.

**Supplemental Figure 8.**
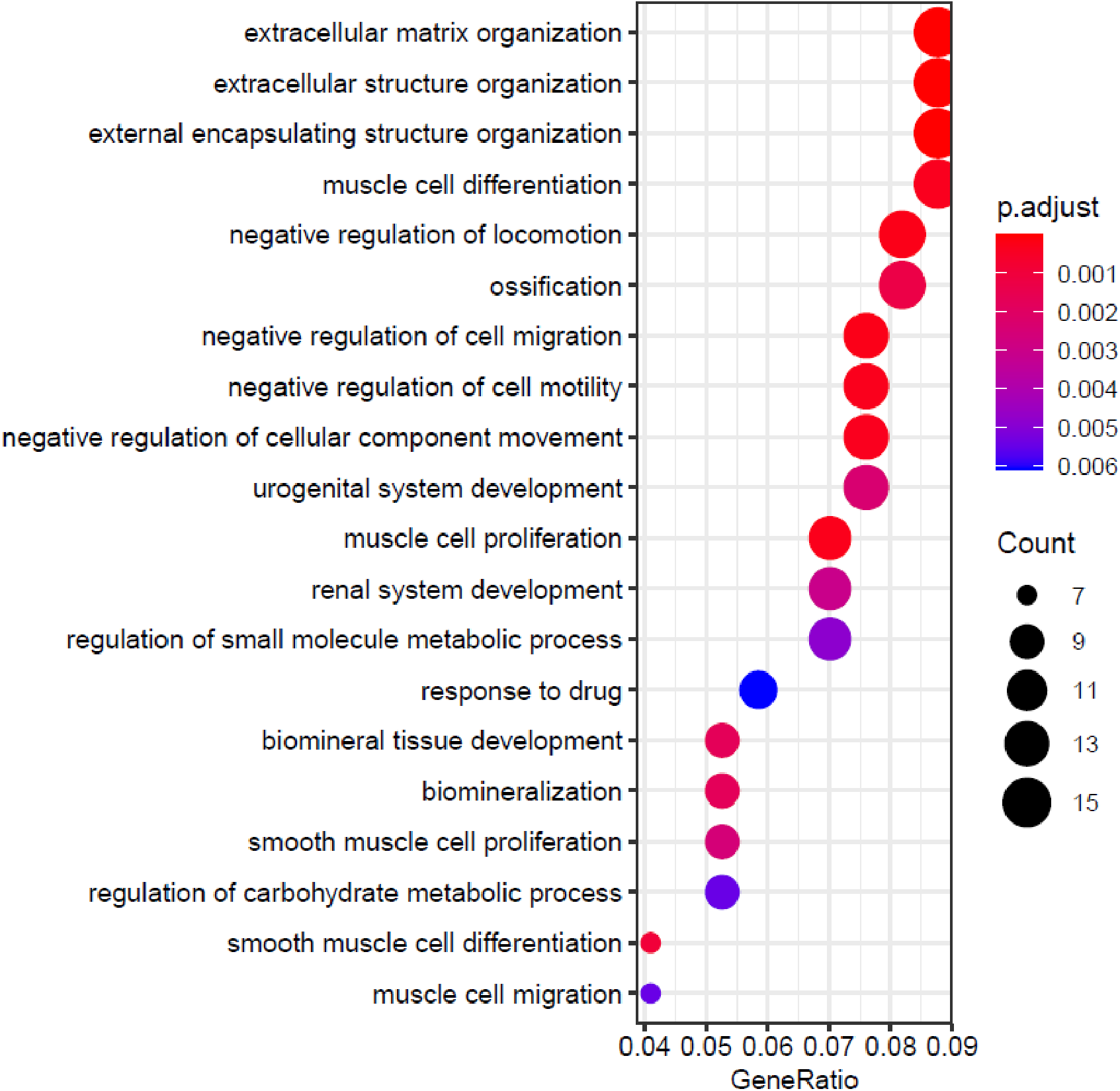
MAF1 overexpression results enrichment for terms related to bone biology. Top 20 biological process related gene ontology (GO) enrichment terms of genes changed by MAF1 overexpression on day 0. Genes with P-adj <0.05 and log2Fold > 0.7 in either direction, were used for analysis.

**Supplemental Figure 9.**
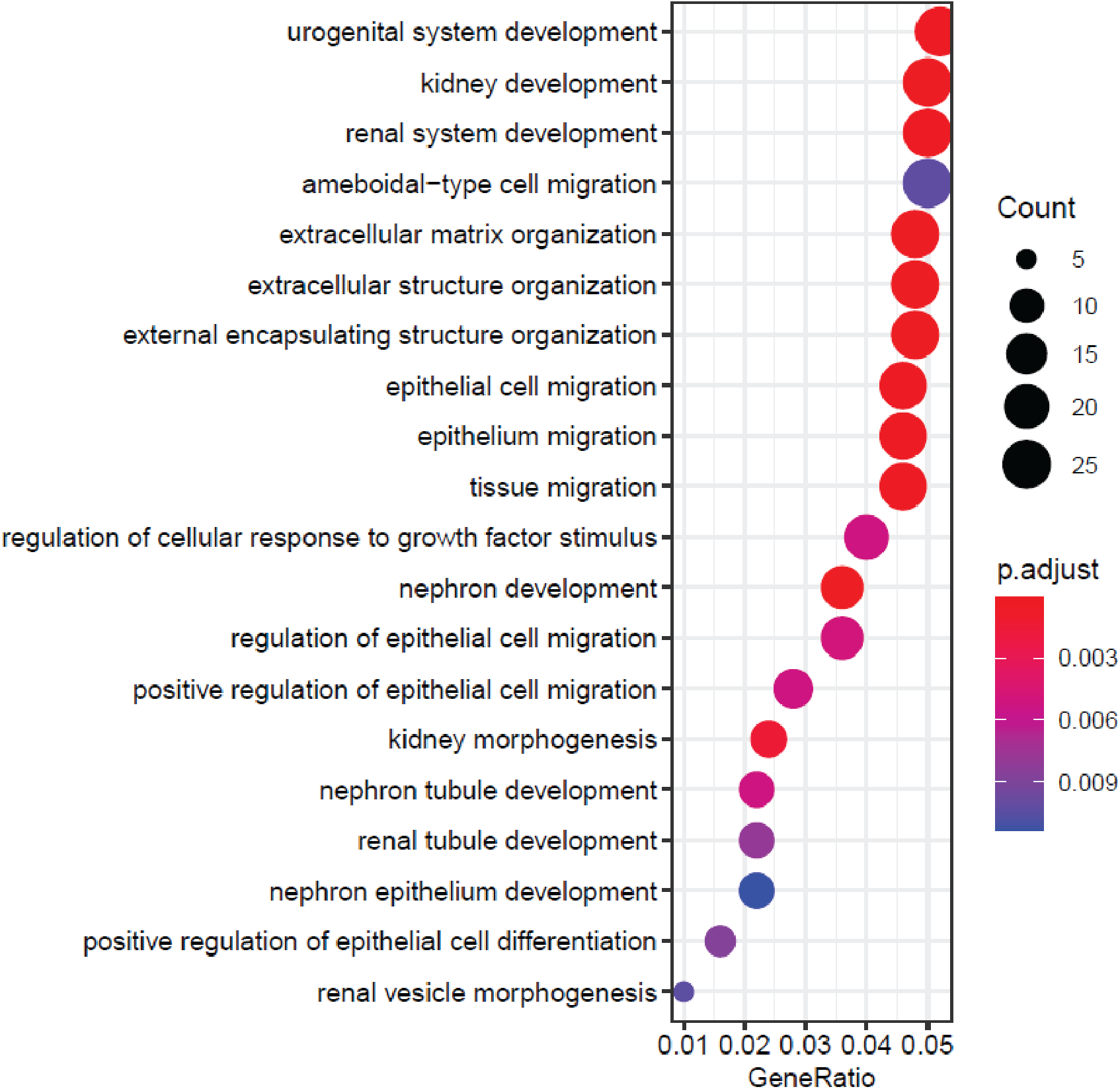
Maf1 knockdown causes enrichment for terms related to bone and renal biology. Top 20 biological process related gene ontology (GO) enrichment terms of genes changed by Maf1 knockdown on day 0. Genes with P-adj <0.05 and log2Fold > 0.7 in either direction, | were used for analysis.

**Supplemental Figure 10.**
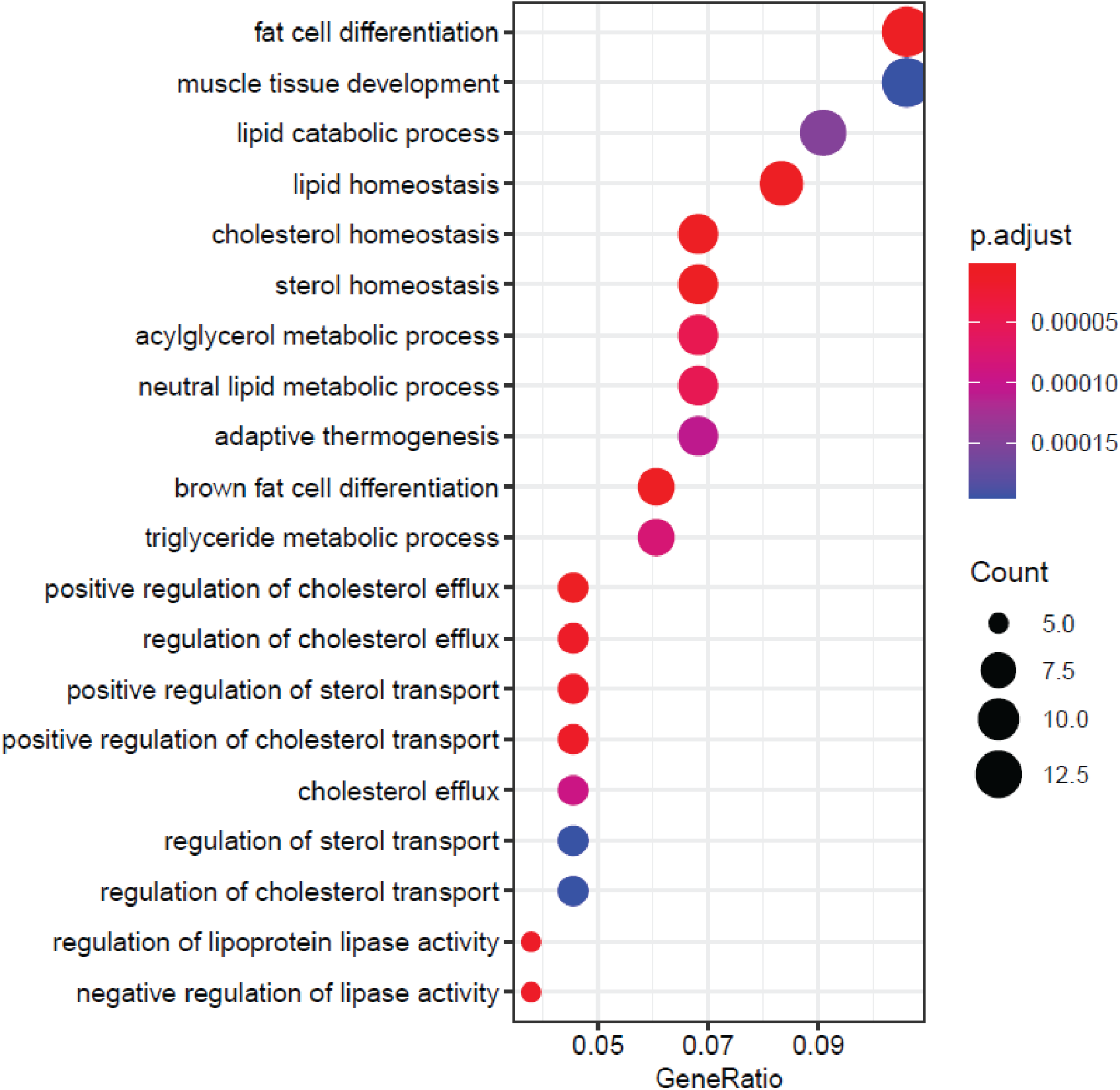
ML-60218 treatment results in enrichment in GO terms related to lipid metabolism. Top 20 biological process related gene ontology (GO) enrichment terms of genes changed by ML-60218 treatment on day 0. Genes with P-adj <0.05 and log2Fold > 0.7 in either direction, were used for analysis.

**Supplemental Figure 11.**
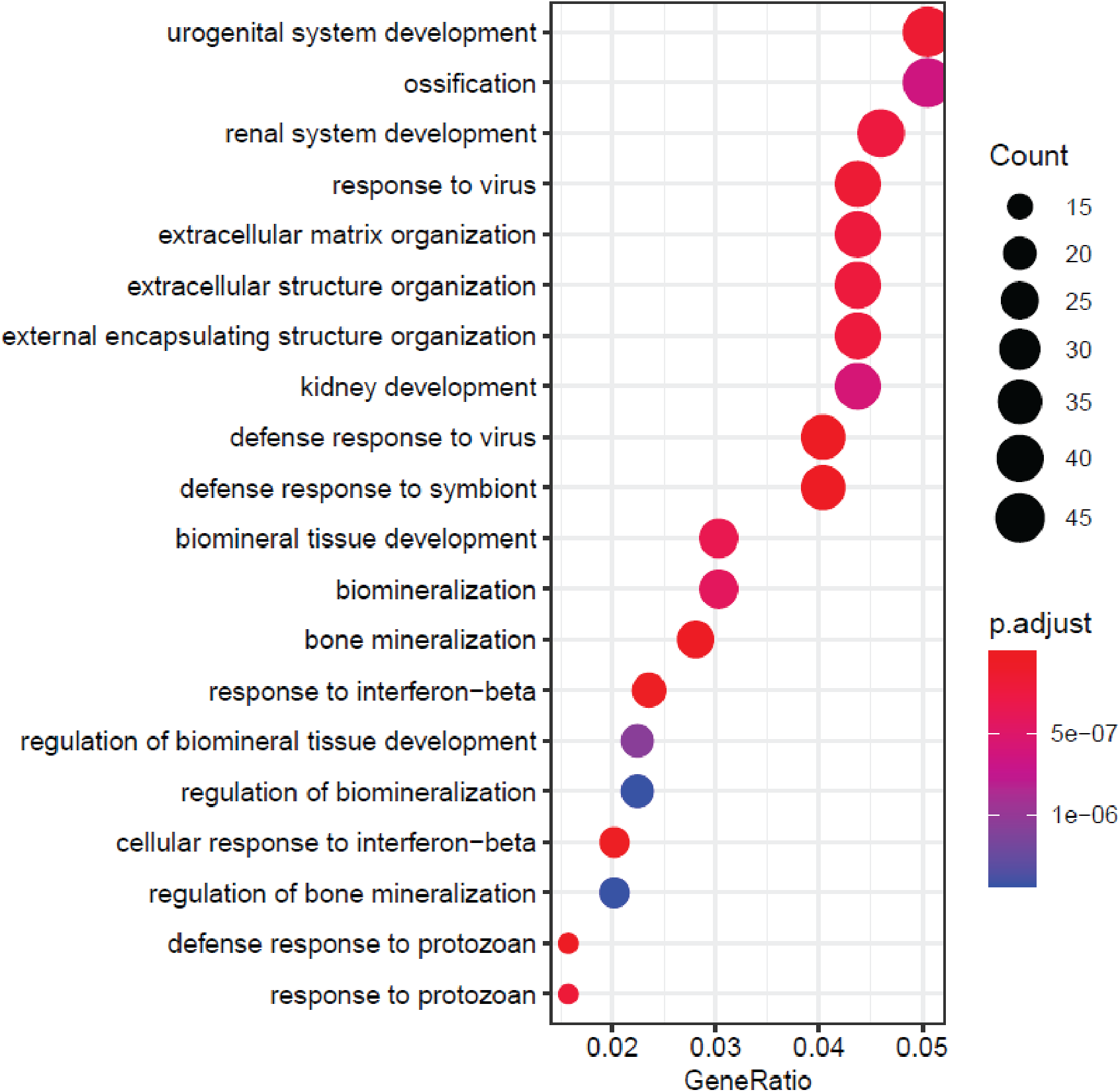
Brf1 knockdown produces gene changes that are enriched in GO terms related to bone biology and immune responses. Top 20 biological process related gene ontology (GO) enrichment terms of genes changed by Brf1 knockdown on day 0. Genes with P-adj <0.05 and log2Fold > 0.7 in either direction, were used for analysis.

**Supplemental Figure 12.**
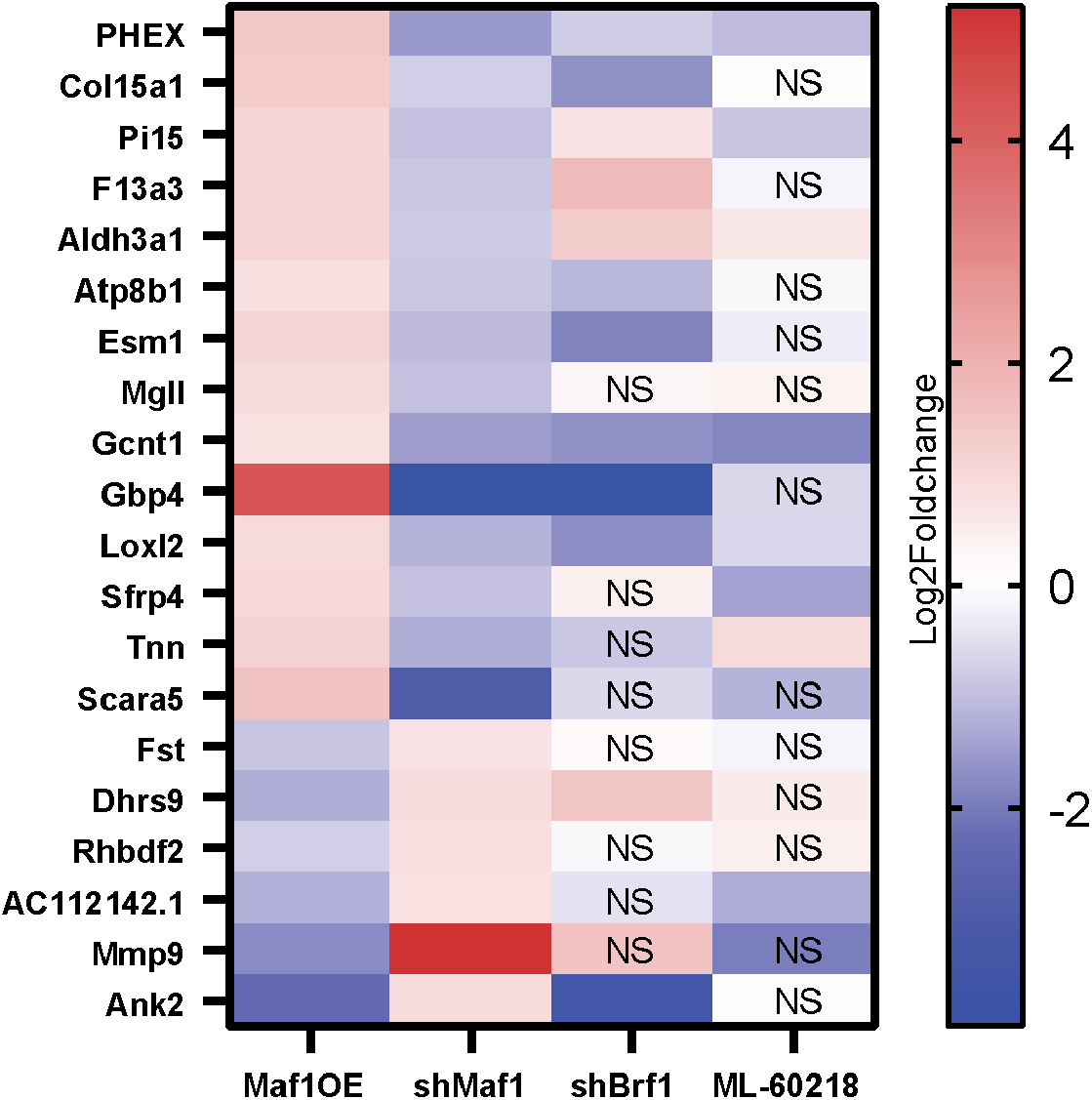
Genes altered by changes in Maf1 expression prior to differentiation. Genes that were significantly altered at day 0 in opposing directions by Maf1 overexpression and Maf1 knockdown by at least log2fold 0.7 are shown. Changes in corresponding genes after Brf1 knockdown or ML-60218 treatment are shown. NS: not significantly affected.

## Tables

**Supplementary table 1.**
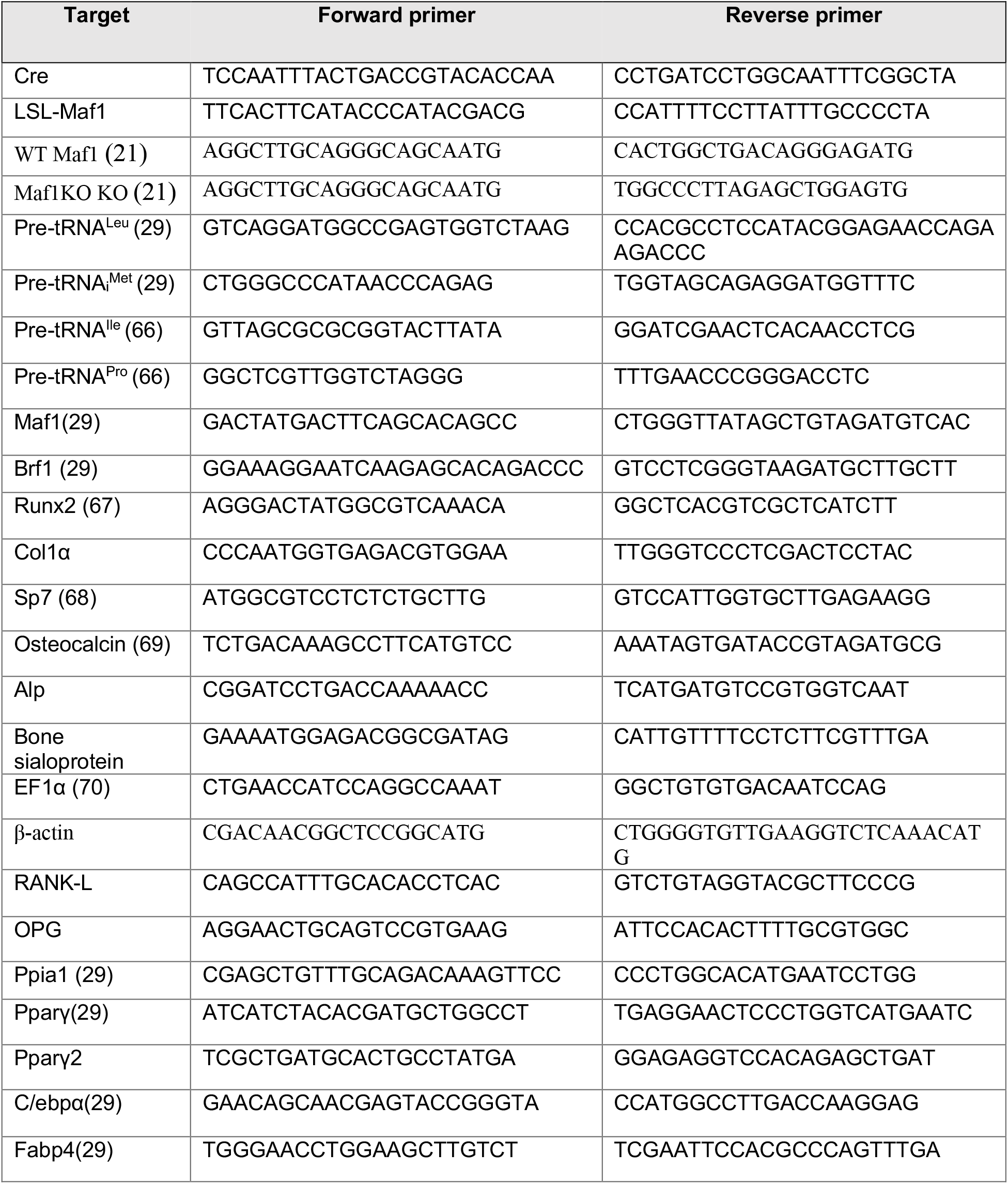
qPCR primers used for genotyping and qRT-PCR analysis.

